# A Theory of Centriole Duplication Based on Self-Organized Spatial Pattern Formation

**DOI:** 10.1101/424754

**Authors:** Daisuke Takao, Shohei Yamamoto, Daiju Kitagawa

## Abstract

In each cell cycle, centrioles are duplicated to produce a single copy of each pre-existing centriole. At the onset of centriole duplication, the master regulator Polo-like kinase 4 (Plk4) undergoes a dynamic change in its spatial pattern on the periphery of the pre-existing centriole, forming a single duplication site. However, the significance and mechanisms of this pattern transition remain largely unknown. Using super-resolution imaging, we found that centriolar Plk4 exhibits periodic discrete patterns resembling pearl necklaces, frequently with single prominent foci. We constructed mathematical models that simulated the pattern formation of Plk4 to gain insight into the discrete ring patterns. The simulations incorporating the self-organization properties of Plk4 successfully generated the experimentally observed patterns. We therefore propose that the self-patterning of Plk4 is crucial for the regulation of centriole duplication. These results, defining the mechanisms of self-organized regulation, provide a fundamental principle for understanding centriole duplication.

## INTRODUCTION

Centrosomes serve as major microtubule-organizing centers, and thus play a fundamental role in animal cells. Each centrosome comprises two centrioles, the microtubule-based core architecture. Centrioles are duplicated once in every cell cycle and only a single copy is generated of each pre-existing centriole in each duplication (Banterle and Gönczy, 2017; Nigg and Holland, 2018). This strict regulation of the number of centriole copies ensures bipolar mitotic spindle assembly and thus optimal regulation of the cell cycle. However, the mechanisms underlying the regulation of the number of centriole copies are largely unknown. In human cells, three centriolar proteins, namely Polo-like kinase 4 (Plk4), STIL, and HsSAS6, have been identified as the core components involved in coordinating the onset of centriole duplication (Banterle and Gönczy, 2017; Nigg and Holland, 2018). In the early G1 phase, Plk4 localizes in a biased ring-like pattern surrounding the proximal periphery of the mother centriole (Ohta et al., 2014, 2018), using CEP152 as a scaffold (Cizmecioglu et al., 2010; Dzhindzhev et al., 2010; Hatch et al., 2010). Following the centriolar recruitment of STIL and HsSAS6, the ring-like pattern of Plk4 changes dynamically into a single focus containing the STIL–HsSAS6 complex (Arquint et al., 2015; Ohta et al., 2014, 2018). Positive and negative regulation, which is based on the bimodal binding mode between Plk4 and STIL, mediates the local restriction of Plk4 at the procentriole assembly site. Although the Plk4 focus exclusively provides the site for cartwheel assembly and subsequent procentriole formation, the significance and molecular mechanisms underlying the dynamic pattern transition of Plk4 are yet to be addressed.

A recent study investigating the molecular dynamics of Plk4 revealed that it possesses intrinsic self-organization properties, such as self-assembly and the promotion of dissociation/degradation of neighboring Plk4 molecules in an autophosphorylation-dependent manner (Yamamoto and Kitagawa, 2018). This finding suggests that: 1) Plk4 itself may generate the bias in the spatial patterns around centrioles, and 2) subsequent loading of STIL and HsSAS6 may reinforce this bias to cooperatively provide the single duplication site. The former point is of particular interest because it may be important in addressing the significance of the pattern transition of Plk4. Considering that Plk4 is the master regulator of centriole duplication (Bettencourt-Dias et al., 2005; Habedanck et al., 2005), self-organized generation of bias may play an important role in the onset of the duplication process. Given the self-organization properties of Plk4, it is reasonable to hypothesize the existence of periodic heterogeneity or, more specifically, spatial domains within the ring-like localization. Therefore, quantitative imaging at higher spatial resolutions may allow the detailed analysis of the spatial patterns of centriolar Plk4.

Currently, there is a lack of theoretical approaches in the research concerning centriole duplication. Despite recent advances, a plausible and experimentally verifiable theory to fundamentally explain the mechanisms underlying centriole duplication is yet to be developed. For example, the link between the molecular nature of Plk4 and its dynamic pattern transition in centrioles remains unclear. A number of experimental and theoretical studies have demonstrated that biological systems commonly comprise self-organization mechanisms at the cellular and tissue levels (Halatek et al., 2018; Saha et al., 2018; Sych et al., 2018; Wheeler and Hyman, 2018). Thus, it is interesting and important to reveal the implications of the self-patterning of Plk4 in the pericentriolar nanospace for centriole duplication, and consequently for cellular function. Mathematical modeling may be a powerful tool in addressing this question. Simulations using appropriate mathematical models rather than qualitative reasoning may reproduce and even predict experimental data, enabling verification of the theory. Based on previous findings related to the self-organizing properties of Plk4 (Yamamoto and Kitagawa, 2018), we here propose the first theory of centriole duplication mechanisms by integrating experimental data and mathematical modeling.

## RESULTS

### The centriolar localization pattern of Plk4 is dynamic and synchronized with the cell cycle

During centriole duplication, Plk4 forms a ring-like pattern around the mother centriole and subsequently, this pattern changes into a single focus, which provides the single duplication site (Arquint et al., 2015; Ohta et al., 2014, 2018). To quantitatively confirm this observation during the actual process of the cell cycle, we monitored centriolar Plk4 dynamics in live cells and correlated them with spatial patterns in fixed cells at a higher resolution. HCT116 cells, in which endogenous Plk4 was tagged with mClover, were synchronized via thymidine arrest, then observed via spinning-disc confocal microscopy after thymidine release (Figure 1A and Figure S1A). Most cells reached mitosis approximately 9 h after thymidine release, and within 1 h after mitotic exit, during the putative early G1 phase, accumulation of Plk4 around the centrioles became apparent (Figures 1A and S1). Centriolar Plk4–mClover signals continued to increase for 3–5 h after mitotic exit, and then began to decrease as the cells progressed toward the next mitosis (Figure 1A).

**Figure 1.**
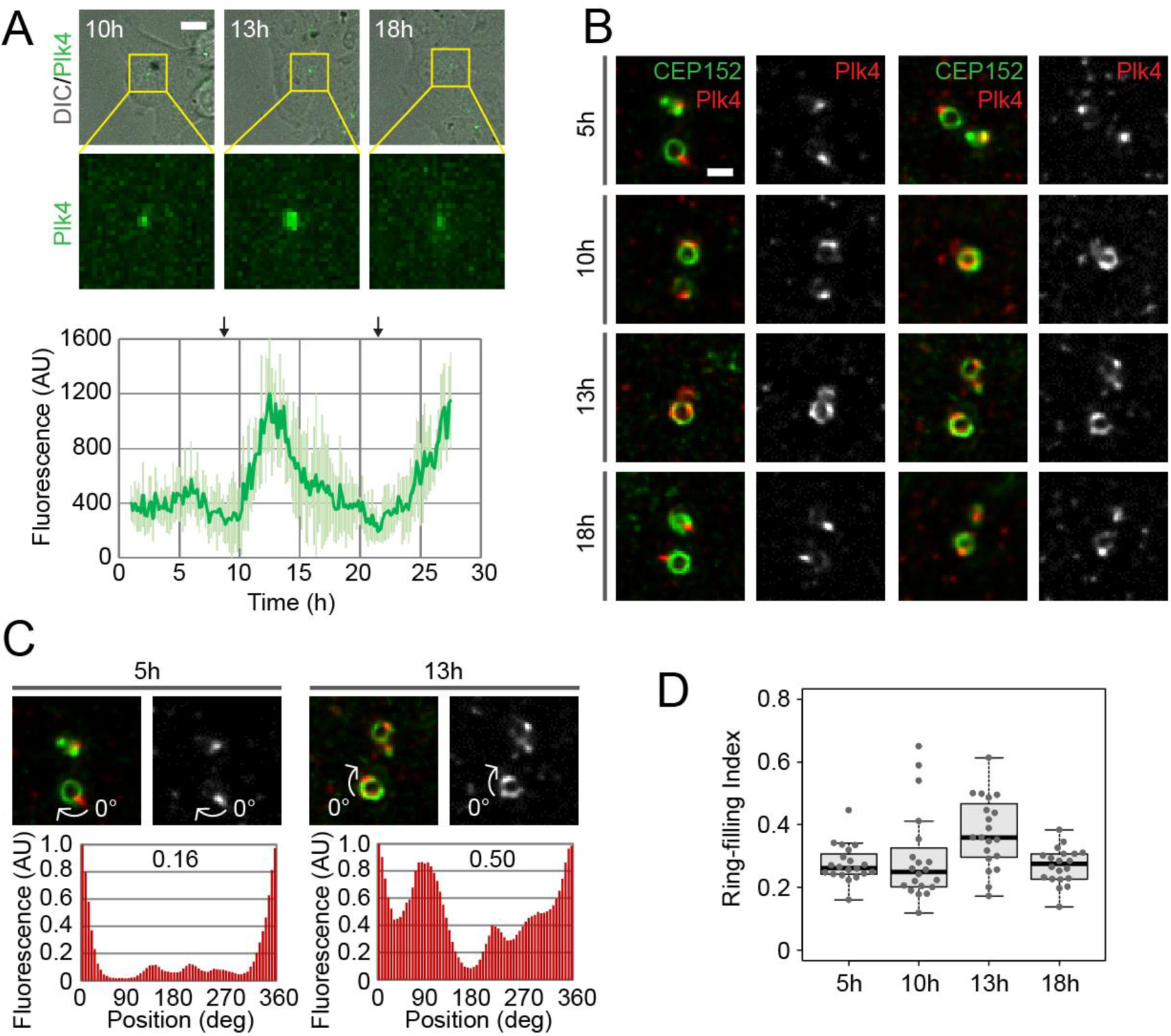
Dynamic changes in the localization pattern of Plk4 during centriole duplication. (A) Live imaging of Plk4 endogenously tagged with mClover in HCT116 cells. Cells synchronized with thymidine were observed at 10 min intervals for 30 h from the time of thymidine release, using spinning-disc confocal microscopy. The graph shows the mean ± SD fluorescence of Plk4–mClover (n = 5 cells). The arrows above the graph indicate approximate time points of metaphase. Scale bar, 10 μm. (B) Confocal immunofluorescence images of endogenous Plk4, with deconvolution. Synchronized cells were fixed at the indicated time after thymidine release and stained with antibodies against Plk4 and CEP152 (centriolar marker). Two representative images are shown for each time point. Scale bar, 0.5 μm. (C, D) Quantification of the Plk4 patterns. Representative oval profiles of Plk4 and the associated ring-filling indices (C), and a plot of all ring-filling indices (D) are shown. Representative images in (C) are from (B). See also Figure S1.

Given the timing of the appearance of the Plk4 ring pattern (early G1 phase, prior to the formation of the daughter centriole) and the single-focus pattern (following the entry of STIL and HsSAS6; Ohta et al., 2014, 2018), the increase and subsequent decrease in the Plk4– mClover signal may reflect the formation and disappearance of the ring pattern, respectively. Indeed, when observed at higher resolution via immunofluorescence confocal microscopy combined with deconvolution, the Plk4 ring pattern most frequently appeared 13 h after thymidine release. This corresponds to the time when the peaks in the Plk4–mClover signal were most frequently detected in the live cells (Figures 1A and 1B). At the onset of the increase in the Plk4–mClover signal in the live cells (10 h), only a small proportion of the fixed cells showed the ring pattern at the higher resolution. When the Plk4–mClover signal decreased as the cells progressed toward mitosis (5 or 18 h), the single-focus pattern became more dominant.

Interestingly, through careful observation, we found that the Plk4 rings in the putative early G1 phase were frequently incomplete. Some resembled Landolt rings (Figure 1B, 10 and 13 h, right panels), while others resembled partial arcs (Figure 1B, 10 h, left panel). Even in the single-focus pattern, the Plk4 sometimes localized weakly in rings (Figure 1B, 18 h). Therefore, instead of merely classifying the patterns into ring or single-focus, we defined a parameter that describes the degree of ring formation and termed it the “ring-filling index”. Using CEP152 as a marker, we determined the ring-filling indices based on normalized oval fluorescence intensity profiles of Plk4 (Figure 1C). The index for a complete ring was 1, and if 50% of the circumference around the centriole was filled with Plk4, the ring-filling index was 0.5. We confirmed that these indices reached their peak 13 h after thymidine release and subsequently decreased (Figure 1D). Combined, these findings demonstrate in a quantitative manner that Plk4 changes its spatial patterns around centrioles dynamically, in synchronization with the cell cycle.

### STED super-resolution microscopy revealed discrete and periodic patterns of centriolar Plk4 localization

We used stimulated emission depletion (STED) super-resolution microscopy to further investigate the incomplete ring patterns of Plk4 at higher resolution. In this analysis, U2OS cells were used instead of HCT116 cells, because they provided better immunofluorescence images and there were no obvious differences in the analyzed data (data not shown). While conventional confocal microscopy with deconvolution allowed us to identify the ring patterns, the resolution was not sufficient to analyze the uneven and incomplete ring patterns in detail (Figures 1B, 2C, and 2A). STED microscopy in combination with deconvolution dramatically improved the spatial resolution, and to our surprise we detected discontinuity in the ring patterns (Figure 2A). Instead of the expected continuous ring, we observed a chain of foci forming a discrete ring (resembling a pearl necklace; Figure 2A).

**Figure 2.**
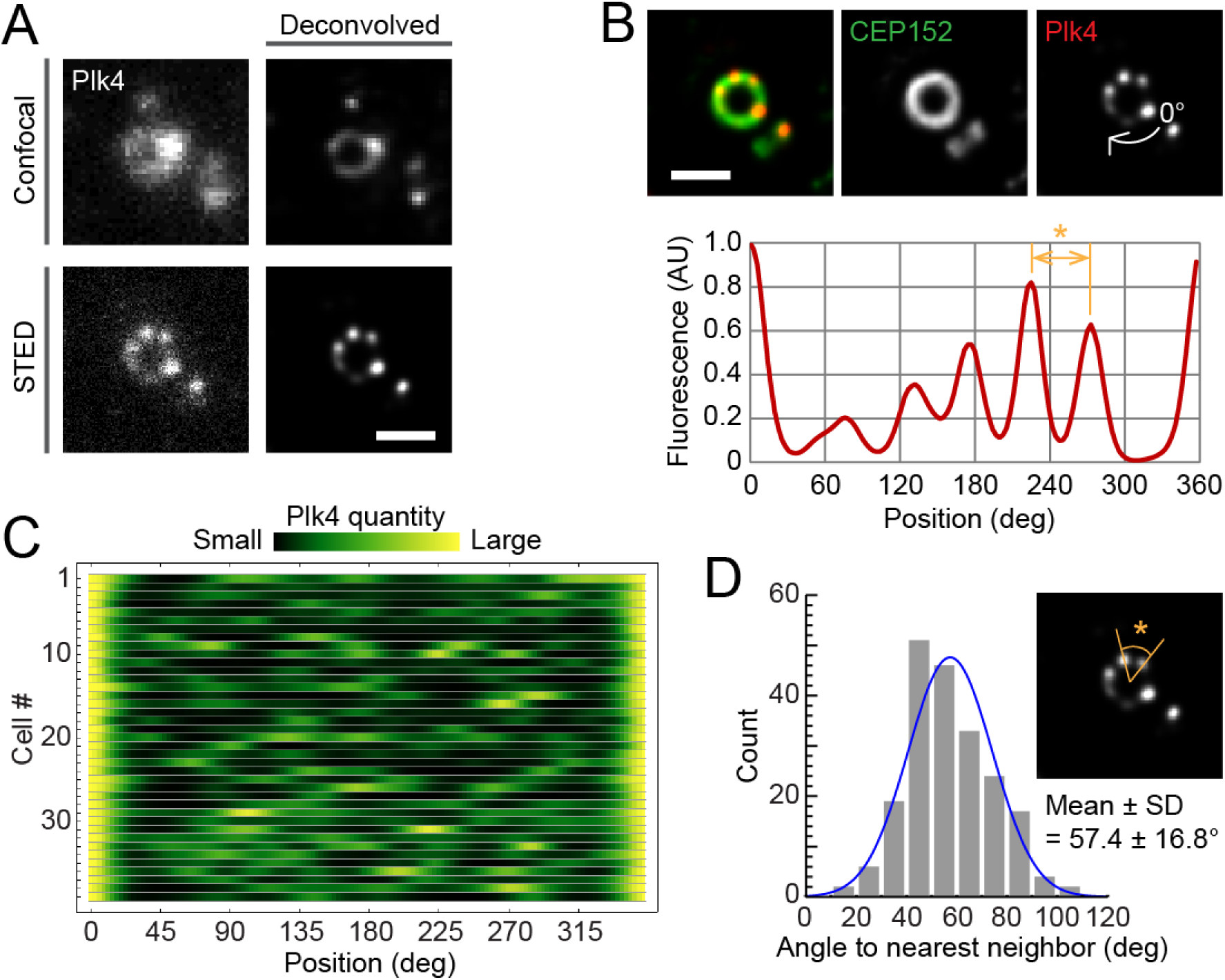
Discrete and periodic Plk4 ring patterns at the super-resolution scale. (A) Comparison of the resolution of conventional confocal and STED images of centriolar Plk4 (from different samples) with or without deconvolution. Scale bar, 0.5 μm. (B) The representative STED image shown in the lower half of (A) merged with CEP152 and the oval profile. The same image is used in all three figures. Scale bar, 0.5 μm. (C) An array plot of all STED profiles of Plk4 (n = 37 cells). (D) Histogram of angles between nearest-neighbor pairs of Plk4 foci in discrete ring patterns. The angle shown in the inset STED image (*) corresponds to the distance indicated in yellow (*) in the profile in (B). See also Figure S2 and S6.

We then extracted oval fluorescence intensity profiles (Figure 2B) to quantitatively analyze the Plk4 ring patterns, similarly to those shown in Figure 1C. The oval profiles we obtained are shown together in Figure 2C as a two-dimensional array plot. In line with previous reports (Ohta et al., 2014, 2018), in most cases we found uneven distribution patterns with single prominent peaks. In addition, as illustrated in Figure 2B, the profiles exhibited spatial periodicity. We detected the peaks of the profiles and measured the angle (the distance in the profile plots) from each peak to its nearest neighbor to quantify the periodicity of the discrete ring patterns. Similar methods have been used successfully to analyze protein localization patterns in the ciliary transition zone or centriole distal appendages (Jana et al., 2018; Shi et al., 2017; Yang et al., 2018). The average angles between nearest-neighbor pairs were approximately 60° (Figure 2D). In other words, the discrete ring patterns tended to show a six-fold rotational symmetry. An alternative method, based on autocorrelation analysis, similarly demonstrated that the period of the discrete pattern was 65.0 ± 16.4° (Figure S2A). This was surprising, given the nine-fold rotational symmetry of the centriole core architecture and the triplet microtubules. Although 40–50° was the most common angle range (Figure 2D), which suggests heterogeneity of the distribution, a 40° periodicity cannot fully explain the distribution, and the majority of angles appeared to be approximately 60°. Therefore, an interpretation other than that based on the nine-fold symmetry of the centriole architecture is required to explain the periodicity of the discrete ring patterns of Plk4.

### The scaffold of Plk4 potentially has double its rotational symmetry and number of slots

We applied the super-resolution analysis to CEP152, the main scaffold of centriolar Plk4, to identify the factor generating the six-fold symmetry of the discrete ring patterns of Plk4. Via STED observations using an antibody to the N-terminal region of CEP152, we found that CEP152 also showed periodic patterns (Figure 3). The rings of CEP152 seemed more continuous than those of Plk4, however, and the periodicities were less obvious than the discrete patterns of Plk4 (Figure 2). Unlike with Plk4, we did not find obvious bias in the periodic patterns of CEP152, which was consistent with previous observations (Ohta et al., 2018). Interestingly, the patterns of CEP152 showed higher spatial frequencies than those of Plk4. The average angle between nearest-neighbor peaks was approximately 30° (Figure 3). CEP152 therefore exhibits a rotational symmetry of order 12, which is double that of Plk4.

**Figure 3.**
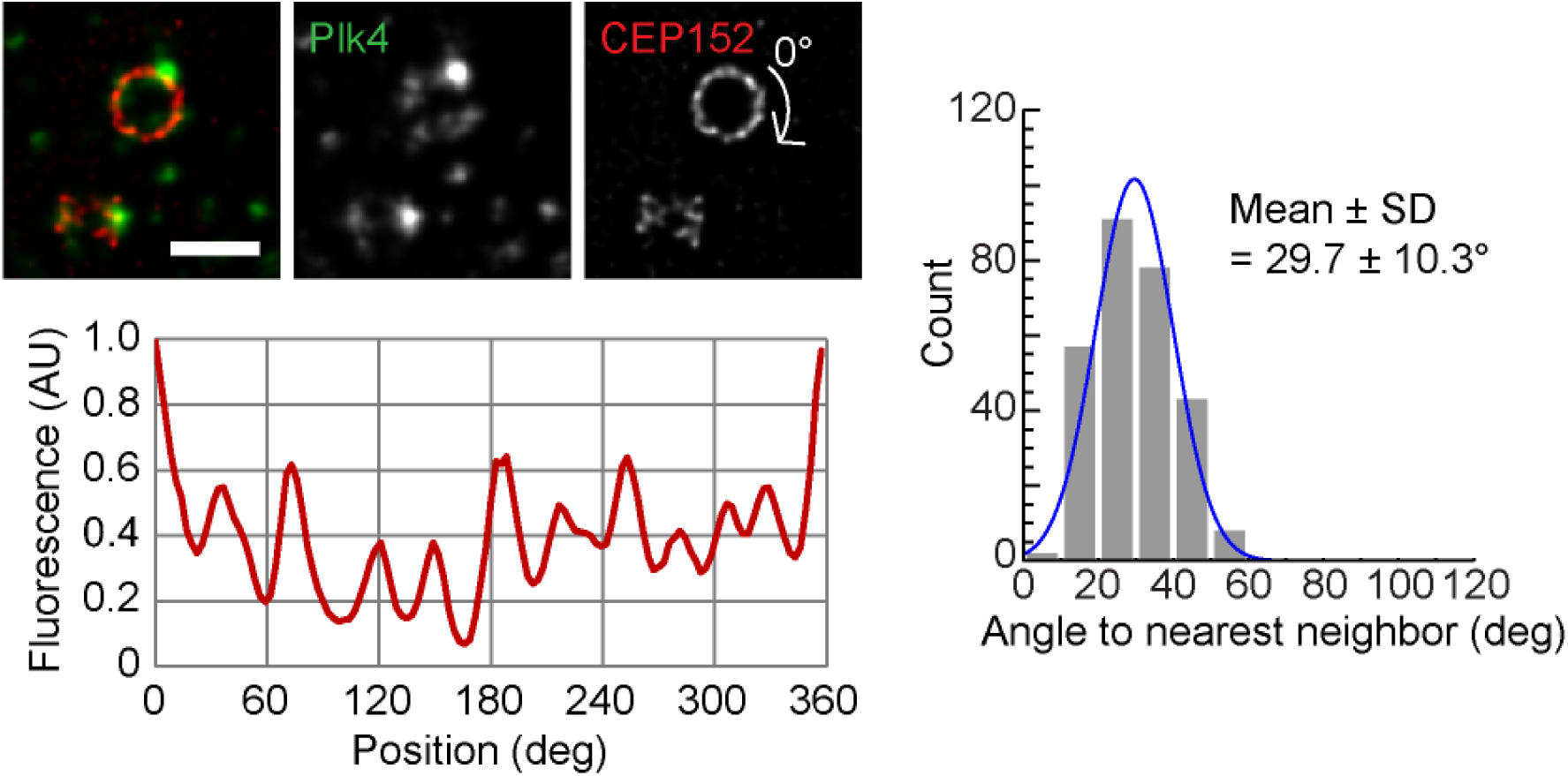
Periodic patterns of CEP152. Representative STED images and quantification of CEP152 spatial patterns. Details as for Figures 2B and 2D. n = 30 cells. Scale bar, 0.5 μm. See also Figure S2 and S6.

Since the periodicity of the CEP152 localization patterns was not entirely clear, we conducted a further analysis for verification. Although on average the discrete patterns of Plk4 exhibited a six-fold symmetry, neighboring Plk4 clusters occasionally localized with approximately 30° intervals between peaks (Figure S2B). This suggests that there may be 12 slots for Plk4 foci. In addition, nearest-neighbor clustering analyses using an antibody recognizing another region of CEP152 (CEP152-Mid) or HCT116 cells confirmed the 30° periodicity or 12-fold rotational symmetry of the CEP152 patterns (Figure S2C). Combined, these results suggest that the scaffold CEP152 provides 12 slots for Plk4, but that Plk4 appears to preferentially select six of them.

### Simulation models based on the self-organization of Plk4 via a lateral inhibition effect reproduce its observed spatial patterns around centrioles

We then investigated how the discrete ring patterns of Plk4, with six uneven foci, are generated from the 12 slots of CEP152. To address this, we reasoned that the molecular nature of Plk4, which has recently been revealed (Yamamoto and Kitagawa, 2018), may provide a clue. Firstly, Plk4 self-assembles into a macromolecular complex (or protein aggregate). Secondly, its properties can switch, depending on its autophosphorylation state. Of note, the phosphorylated (“active”) form of Plk4 is more dynamic in dissociating from centrioles than the non-phosphorylated (“inactive”) form. In addition, the active form of Plk4 promotes the dissociation of inactive Plk4 through self-activation (trans-autophosphorylation) feedback. These properties of Plk4 suggest that it may form the discrete ring patterns by excluding neighboring Plk4 molecules in an autophosphorylation-dependent manner, whereas existing Plk4 complexes gain their mass through self-assembly. This mutual relationship between the active and inactive forms of Plk4 is similar to that of the reaction–diffusion system (also known as the Turing model) or its analogue, the lateral inhibition system, which forms periodic patterns at the cellular or tissue level (Barad et al., 2011; Kondo and Miura, 2010; Liao and Oates, 2017; Nakamura et al., 2006). Accordingly, here we propose the first verifiable theory explaining the mechanism by which the self-organization properties of Plk4 form dynamic spatial patterns and exclusively provide the single duplication site during centriole duplication. The core concept of this theory, which is based on a recently proposed speculative model (Yamamoto and Kitagawa, 2018), is shown in Figure 4.

**Figure 4.**
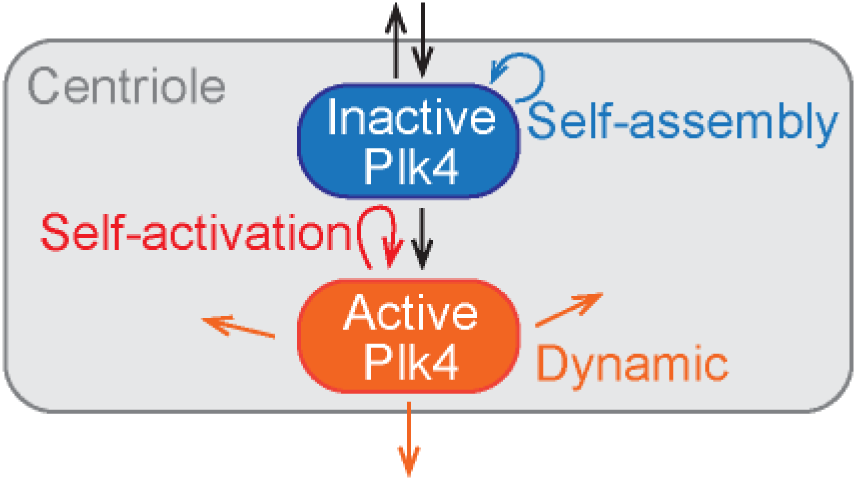
The core concept of the theoretical model. Our theory assumes the presence of two distinct forms of Plk4: inactive and active. While the inactive form interacts with nearby molecules to self-assemble and is activated through autophosphorylation, the active form trans-autophosphorylates adjacent Plk4 molecules (self-activation). The active form is more dynamic (mobile) within and outward the centriole. The self-activation feedback promotes the dissociation of Plk4 from the centriole. The dissociation of Plk4 is attenuated when either form reaches the threshold quantity for the formation of an aggregate complex. These properties of Plk4 result in its lateral-inhibition self-patterning behavior.

Our theory assumes that the active form of Plk4 is dynamic and mobile within the periphery of centrioles. These properties render the active Plk4 capable of reaching and interacting with other distant Plk4 molecules along the centriole periphery. This can cause a lateral inhibition effect, in which active Plk4 molecules actively repel nearby inactive Plk4 molecules by promoting activation and subsequent dissociation. Assuming that this lateral inhibition effect exists, we constructed a mathematical model to simulate the pattern formation of Plk4 (Figure 5A). Given the periodicity of CEP152 scaffolds (Figure 3), we placed 12 segments as Plk4 slots around the centriole. The last segment (*x* = 12) was connected to the first (*x* = 1) to form a closed-loop centriolar ring (Figure 5A). As described above, Plk4 can take two distinct forms: active and inactive. The inactive form influxes from the cytosol (with the kinetic constant *k*1) to the centriolar segments, and both forms dissociate from the centriole (with the distinct kinetic constants *k*2 and *k*5, respectively). The active form was set to be more dynamic in its dissociation (*k*5 > *k*2) and mobile in its interaction with other Plk4 molecules in the neighboring segments (bold orange arrows in Figure 5A). The inactive form self-assembles (*k*6) to cause a positive-feedback effect and turns into the active form through autophosphorylation (*k*3). The self-assembly rate is proportional to the concentration of Plk4 in each segment and in the cytosol. Therefore, all Plk4 influx into each segment was included in the self-assembly term. The active form of Plk4 promotes the activation of adjacent inactive Plk4 in its own segment and in neighboring segments (*k*4). This promotes the dissociation of activated Plk4, generating the lateral inhibition effect. Importantly, when the total amount of Plk4 in a segment reaches a given threshold, the molecules are assumed to form a stable complex (aggregate) (Yamamoto and Kitagawa, 2018), which reduces the dissociation rates of both forms of Plk4. A random fluctuation (±10%) was added to the self-assembly term, so the results differ in each simulation. The initial seeds (input) were provided as random real numbers between 0 and 1. It should be noted that the model does not distinguish between dissociation and degradation, so the observed dissociation may be partly degradation. The units for the kinetic constants and time are arbitrary.

**Figure 5.**
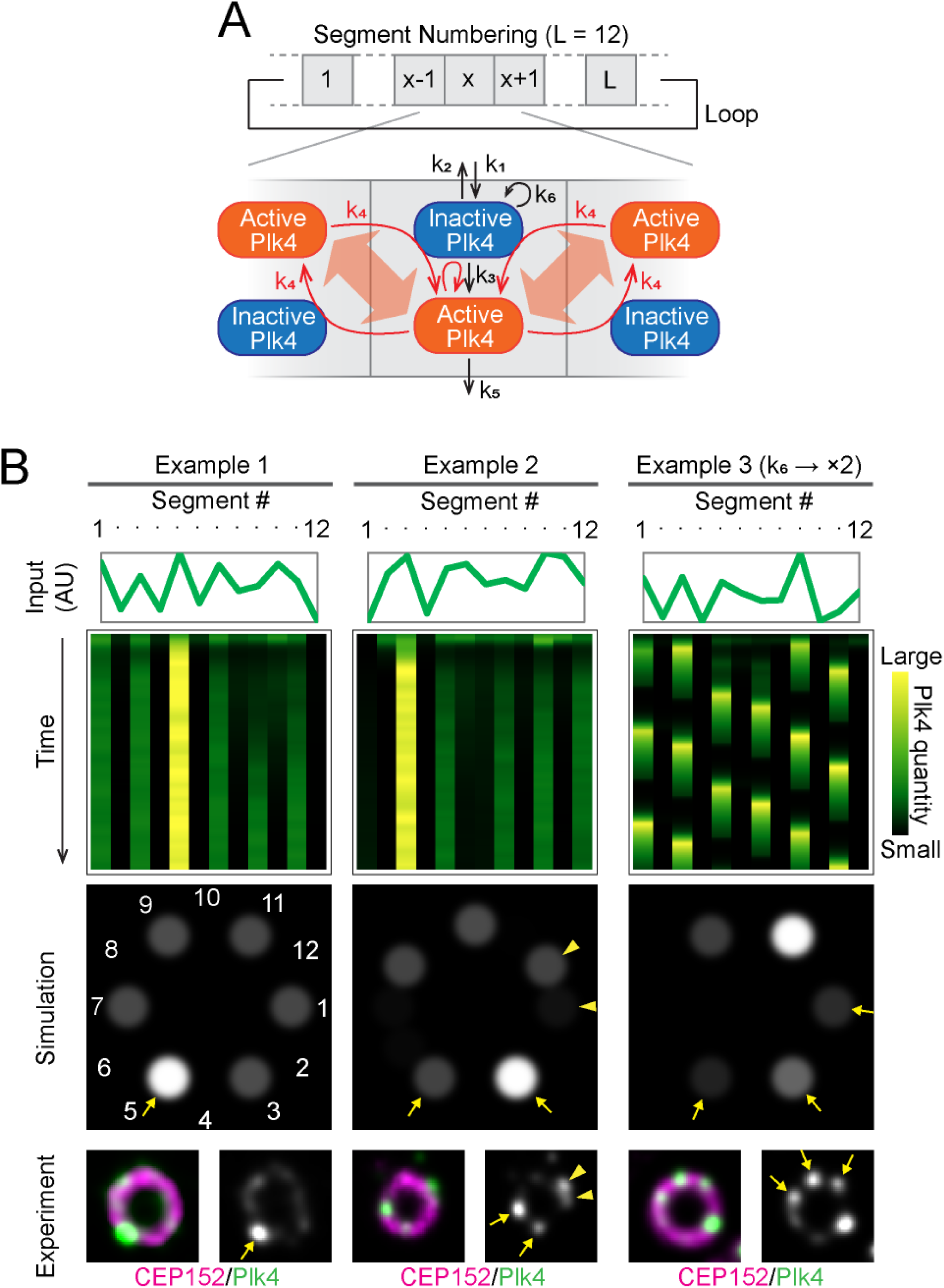
Simulations of the pattern formation of Plk4 based on the lateral inhibition model. (A) Schematic drawing of the lateral inhibition model. (B) Examples of simulation results. Array plots showing the time course of pattern formation, with the initial quantities of Plk4 indicated in the line graphs above (top), and the spatial pattern at the last time point (middle), are shown for each example. Representative STED images resembling the simulation results are shown at the bottom. The arrows and arrowheads indicate the similarities in the patterns between the simulation and the actual observation. The basic parameter setting was used for Examples 1 and 2, but *k*6 was doubled for Example 3. See also Figure S3 and S4, and the STAR Methods section.

Initially, we optimized each parameter of the model (Table S1) to reproduce actual observations. Consequently, approximately six foci of Plk4, containing various quantities of the molecule, appeared from the 12 slots (Figure 5B). For simplicity, the representative results illustrated show the total quantities of Plk4 (i.e. both inactive and active), even though each focus typically included a certain proportion of active Plk4. Due to the lateral inhibition effect, Plk4 tended to occupy alternate slots to form the distinct “pearl necklace” pattern seen in the STED images (Figure 5B). Initial random patterns frequently converged into discrete ring patterns as a result of competition between the slots. Notably, most of the discrete ring patterns generated in the simulations had single prominent foci, similarly to those in the STED images. The pattern may be more dynamic, however, depending on the parameters used. For example, when the *k*6 value was twice that used in Examples 1 and 2 (Figure 5B), the spatiotemporal patterning of Plk4 became more dynamic and resulted in a multiple oscillation mode (Example 3). Nevertheless, it still exhibited discrete patterns with a prominent peak at each time point. This variation in the spatial and temporal patterns simulated may explain the variety of the spatial patterns of Plk4 observed in our STED analysis. The lateral inhibition model proposed in this study thus reproduces the observed Plk4 patterns at centrioles well.

The model assumes 12 centriolar slots for Plk4, based on the STED analyses (Figures 3, S2B, and S2C). Given the somewhat unclear periodicity of the CEP152 spatial patterns, we tested the robustness of the model with respect to the periodicity of the scaffolds. All simulations using different numbers of slots (9, 10, 11, or 13) resulted in the formation of pearl necklace patterns (Figure S3), as in the simulations using 12 slots (Figure 5). Regardless of the number of slots, therefore, the spatial patterns of Plk4 were similar, albeit with slightly different spatial frequencies. The lateral inhibition effect of Plk4 is thus not strictly dependent on the number of slots, and this model allows flexibility in scaffold conformation.

Next, to test the validity of the core concept that the self-organization properties of Plk4 alone can cause the discrete ring patterns to form, we tested how altering the mobility of Plk4 would affect the model. For this alternative mathematical model, termed the reaction–diffusion model for convenience, we used the core concept shown in Figure 4, but more explicitly considered the lateral diffusion of Plk4 within the periphery of the centriole (Figure S4A). This is possible since the pericentriolar space is filled with a number of proteins and therefore serves as a diffusion trap to tether Plk4 around the centrioles. Similarly to the lateral inhibition model (Figure 5A), the active form of Plk4 was assumed to be more dynamic (i.e., larger diffusion coefficient) than the inactive form (Figure S4A and Table S2). The active form repels the inactive form through self-activation feedback, although only within the same segment (Figure S4A). Simulations using this model also resulted in the formation of discrete ring patterns (Figure S4B). Similarly to the lateral inhibition model (Figure 5B), Plk4 formed uneven foci aligned along the circumference, regardless of the number of segments (Figure S4B). These results further support our theory that the intrinsic properties of Plk4 mean that it can self-organize into the patterns around centrioles (Figure 4). For simplicity, hereafter we adopt and further verify the lateral inhibition model (Figure 5A) as a tentative model.

### The lateral inhibition model reproduces centriole duplication under physiological and perturbed conditions

Using the lateral inhibition model, we then simulated the pattern transition of Plk4 in the presence of STIL and HsSAS6. The ring-like pattern changes into the single-focus mode following the centriolar entry of STIL and HsSAS6 at the onset of procentriole formation (Ohta et al., 2014, 2018). We therefore expected to see this pattern transition in the simulations. The mechanism through which STIL and HsSAS6 cooperatively regulate the pattern transition to restrict the duplication site remains unclear. In the model, therefore, we only included the centriolar entry of STIL, using it to represent the STIL–HsSAS6 complex. The time-evolved simulations were set as follows: 1) initially, the random seeds of Plk4 evolved to form discrete ring patterns via the lateral inhibition effect, as shown in Figure 5B; 2) next, the expression level of STIL began to increase, as observed in the late G1 to S phases in human cells (Arquint and Nigg, 2014; Arquint et al., 2012; Izraeli et al., 1997; Tang et al., 2011); 3) finally, STIL entered the Plk4 slots in which the quantity of active (phosphorylated) Plk4 was above a given threshold (based on reports showing that the centriolar loading of STIL depends on Plk4-mediated phosphorylation; Dzhindzhev et al., 2014, 2017; McLamarrah et al., 2018; Moyer et al., 2015; Ohta et al., 2014). Notably, in this model the centriolar loading of STIL was biased according to the Plk4 bias within the discrete ring pattern. In other words, Plk4–STIL interactions at centrioles were assumed to occur stochastically, as in general protein–protein interactions. The other condition we considered in the simulations was that, once the cytosolic level of STIL had begun to increase, the cytosolic level of Plk4 was assumed to decrease. It has previously been reported that increased quantities of STIL in the cytoplasm may promote the degradation of Plk4 (Arquint et al., 2015; Moyer et al., 2015; Ohta et al., 2018). Thus the assumption that the cytosolic level of Plk4 decreases after the centriolar loading of STIL seems reasonable, despite the lack of biochemical studies specifically investigating the expression levels of endogenous Plk4. In addition, it is also possible that the dissociation/degradation of Plk4 in the slots in the absence of STIL is promoted via negative-feedback regulation, mediated by the Plk4–STIL interaction (Ohta et al., 2018). As expected, simulations including such negative-feedback regulation yielded similar results but with slightly faster transition from the discrete ring into the single focus after centriolar loading of the STIL–HsSAS6 complex (Figure S5).

Consequently, simulations using the lateral inhibition model reproduced the Plk4 pattern transition from the discrete ring into the single focus upon centriolar loading of the STIL–HsSAS6 complex (Figure 6A). The most prominent focus of Plk4 in the rings tended to remain after STIL–HsSAS6 loading. In addition, we simulated a perturbation of centriole duplication through overexpression of the components and compared the simulated results with experimental results to further verify this model. Overexpression of Plk4, STIL, or HsSAS6 is known to induce overduplication of centrioles (Arquint et al., 2012; Habedanck et al., 2005; Kleylein-Sohn et al., 2007; Strnad et al., 2007; Tang et al., 2011; Vulprecht et al., 2012). In simulations using overexpression of Plk4 (the cytosolic level of Plk4 was set five times higher), multiple Plk4 foci tended to remain even after STIL–HsSAS6 loading. This was consistent with the experimental results (Figure 6B). This is due to increased influx of Plk4 into the centriole, resulting in multiple foci retaining Plk4 quantities above the threshold of STIL binding. Despite this increased influx, the lateral inhibition effect remained. Interestingly, under overexpression of Plk4, the simulated Plk4 patterns were relatively unstable throughout the time course (Figure 6B), until they were stabilized via the entry of STIL.

**Figure 6.**
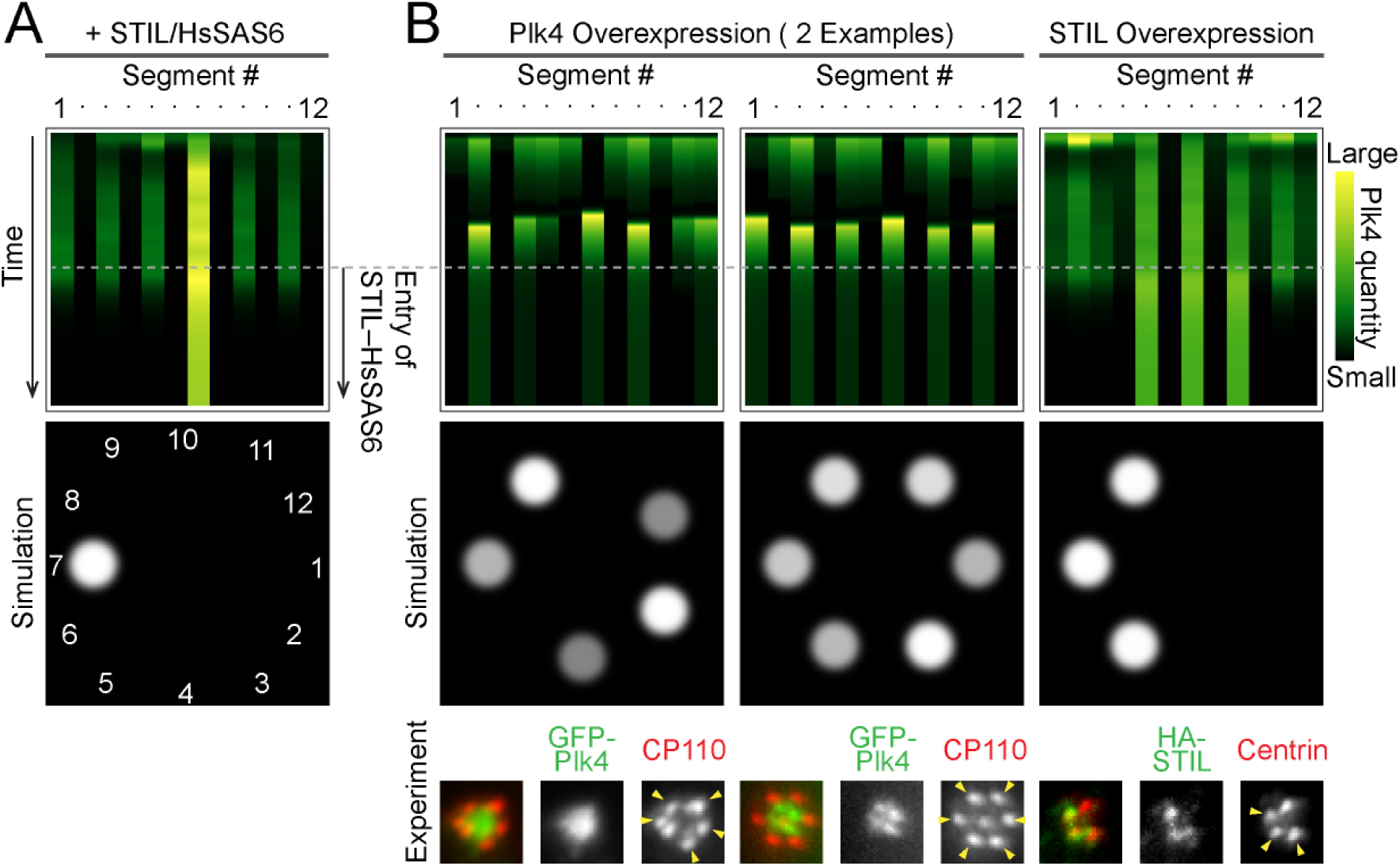
Simulations using centriolar entry of STIL–HsSAS6 and exogenous perturbation. (A) Representative simulation results using the entry of the STIL–HsSAS6 complex. At the mid-point of the time course, the STIL–HsSAS6 complex begins to accumulate. (B) Representative simulation results using overexpression of Plk4 or STIL. Representative experimental data showing similar patterns are shown at the bottom. The arrowheads in the images indicate the duplicated centrioles. See also Figure S5.

Overexpression of STIL was then simulated by setting a lower threshold (one fifth) of STIL binding to Plk4 foci. Under this condition, several Plk4 foci tended to remain after STIL– HsSAS6 loading, which was also consistent with our experimental observations (Figure 6B). It is known that overexpression of HsSAS6 also induces overduplication of centrioles (Strnad et al., 2007). In our model we represented the involvement of HsSAS6 in the form of the STIL– HsSAS6 complex, but we hypothesize that HsSAS6 may stabilize Plk4 foci cooperatively with STIL by forming a trimeric complex. Combining these findings, we conclude that the model successfully reproduced our observations under perturbation via the overexpression of major centriole duplication factors, regardless of physiological conditions. This further supports our hypothesis that the self-organization properties of Plk4 may be one of the mechanisms regulating centriole duplication (Figure 4).

### Inhibition of Plk4 activity resulted in unexpected patterns, implying flexibility in the pattern of the Plk4 scaffold

We also tested the effects of inhibition of the kinase activity of Plk4. Assuming that Plk4 is converted into its active form via autophosphorylation, inhibition of its kinase activity may decelerate the inactive-to-active transition of Plk4, thereby diminishing the lateral inhibition effect (Figure 4). Using lower rates of the inactive-to-active transition, our model predicted that Plk4 would occupy all of the available slots and exhibit 12 foci symmetrically arranged around a centriole (data not shown).

In contrast, our STED observations showed that Plk4 exhibits nine-fold rotationally symmetrical spatial patterns upon inhibition of its kinase activity by treatment with the Plk4 inhibitor centrinone (Figure S6A; Wong et al., 2015). The lateral inhibition model based on 12 slots was unable to explain this pattern. A possible explanation for this discrepancy is that CEP152 (the scaffold for Plk4) may also change its patterns around centrioles following treatment with centrinone. We used STED microscopy to analyze the spatial patterns of CEP152 after treatment with centrinone to test this possibility. Interestingly, we found that, in the presence of centrinone, the middle part of CEP152 exhibited patterns closer to nine-fold symmetry (40° intervals), whereas the N-terminal part continued to hold approximately 12 slots (30° intervals) at the centriole (Figure S6B). This suggests that the behavior of the middle part of CEP152 depends on the kinase activity of Plk4, and may also control the pattern formation of centriolar Plk4. Collectively, these findings suggest that following the inhibition of kinase activity via treatment with centrinone, the mode of interaction between Plk4 and CEP152 changes, and the mechanism of complex formation is no longer within the scope of the lateral inhibition model. However, considering the somewhat unclear periodicity of CEP152, further investigation is warranted to improve our understanding of the effects of treatment with centrinone on the centriolar pattern formation of Plk4.

## DISCUSSION

This is the first study to integrate imaging-based experimental data and mathematical models concerning the self-organization properties of Plk4, and to propose the theory that these properties play a critical role in the regulation of centriole duplication. Firstly, using STED super-resolution microscopy and subsequent quantitative analyses, we revealed that Plk4 forms periodic discrete ring patterns on the periphery of centrioles. We found that the periods of the patterns were on average around 60°. In other words, the discrete ring patterns of Plk4 predominantly have a six-fold rotational symmetry. Moreover, the levels of Plk4 within these spatial patterns tended to be biased towards one of the foci within the ring. On the other hand, similar analyses revealed unbiased 12-fold rotational symmetry in the CEP152 scaffold in its centriolar localization patterns. Secondly, we constructed mathematical models to simulate the pattern formation of Plk4 during centriole duplication, based on recent findings indicating that Plk4 may change its dynamics and localization via its intrinsic properties (Yamamoto and Kitagawa, 2018). The lateral inhibition model we constructed reflected the core concept of our theory: accumulated Plk4 repels neighboring Plk4 molecules via self-activation feedback in the pericentriolar spatial domain (Figures 4 and 5). Indeed, the model reproduced most of our experimental data, with one exception.

As schematically shown in Figure 7, our theory explains the involvement of molecular dynamics in the determination of centriole duplication sites. In this model, centriole duplication proceeds as follows. Plk4 first enters the centriolar scaffold to form randomly-distributed seeds and then competition occurs, via the lateral inhibition effect, which results in the appearance of discrete ring patterns. The model also includes the self-assembly property of Plk4 (Yamamoto and Kitagawa, 2018), by which condensed Plk4 attenuates its dissociation rate. Therefore, the slot that recruits the greatest quantity of Plk4 at the onset is favored, in that it prevents neighboring slots from recruiting Plk4 molecules and tends to survive as the largest Plk4 focus via first-come-first-served and the-rich-grow-richer processes (“Self-patterning” in Figure 7). Through stochastic means, STIL and HsSAS6 preferentially bind to the largest Plk4 focus in the discrete ring, leading to stabilization. This assumption is reasonable, given that stochastic interactions between Plk4 and STIL–HsSAS6 are probably biased such that the more Plk4 there is in a focus, the more frequently the molecules interact with each other. Increasing expression levels of STIL in turn promote the degradation of cytosolic Plk4, resulting in decreased entry of Plk4 into the centriole. Given that the population of phosphorylated Plk4 in centrioles increases after centriolar loading of the STIL–HsSAS6 complex (Ohta et al., 2018; Yamamoto and Kitagawa, 2018), it is also conceivable that the negative-feedback regulation based on the Plk4– STIL interaction promotes the dissociation/degradation of Plk4 around the mother centriole wall, except at the duplication site (Ohta et al., 2018). Consequently, the largest Plk4 focus remains as the only site of centriole duplication.

**Figure 7.**
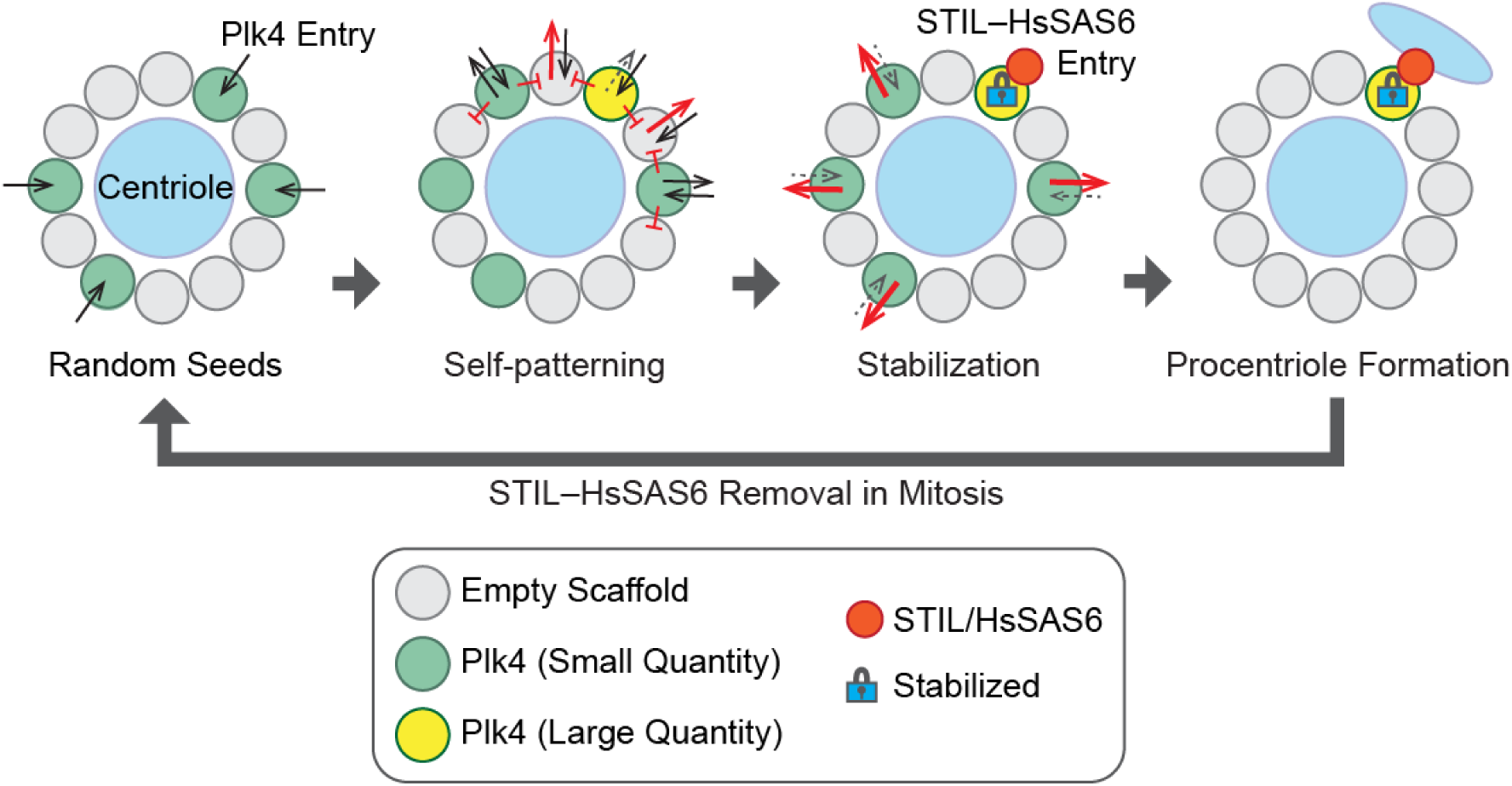
Model of centriole duplication. The self-patterning theory explains how the self-organization properties of Plk4 result in the formation of the biased discrete ring patterns to provide the single centriole duplication sites.

According to our self-patterning theory, Plk4 can independently form periodic discrete patterns, i.e., potential duplication sites. Most importantly, because of its intrinsic properties, Plk4 alone can generate bias from random seeds in the periphery of pre-existing centrioles. However, the self-patterning of Plk4 exhibits fluctuation and plasticity until it is stabilized by STIL and HsSAS6. This two-step process may serve as backup and buffer mechanisms to ensure the precise regulation of centriole duplication. Indeed, in biological systems, it is commonly observed that initial symmetry-breaking steps induce weak bias, and subsequent feedback mechanisms robustly fix the asymmetry (Chen et al., 2018; Goryachev and Leda, 2017; Kim et al., 2018). Instead of forming the definitive duplication site from the beginning, the weakly biased ring patterns of Plk4 may provide backup sites, while the entry of STIL– HsSAS6 ensures that only one site is determined for centriole duplication. In this regard, it is interesting that the spatiotemporal patterning of Plk4 may be more dynamic, depending on the parameters used in simulations (Figure 5B, Example 3). Thus, our theory predicts the possibility that potential centriole duplication sites switch dynamically within the periphery of centrioles. However, such a system may not be sufficiently stable to ensure the formation of the single duplication sites. The development of future techniques enabling live-cell super-resolution imaging may provide the answer.

Our simulations including the overexpression of Plk4 or STIL both resulted in centriole overduplication, as actually observed in our experiments (Figure 6B). Multiple Plk4 foci were stabilized to provide centriole duplication sites. Interestingly, the numbers of stabilized Plk4 foci in the overexpression simulations did not exceed six, potentially explaining the limitation in the maximum number of overduplicated daughter centrioles. Indeed, most previous studies have convincingly demonstrated that six is the maximum number of daughter centrioles that can be formed on a single mother centriole (Arquint et al., 2012; Habedanck et al., 2005; Kleylein-Sohn et al., 2007; Strnad et al., 2007; Tang et al., 2011; Vulprecht et al., 2012). However, it is also possible that this may be merely because of space limitations. Importantly, we provide the first theoretical explanation and prediction of the mechanisms involved in centriole overduplication.

It is significant that this first theory for centriole duplication is based on both precise observations and mathematical modeling. However, further experimental evidence and improvement of the present models is required to elucidate these processes. For example, actual parameters for molecular dynamics, such as association/dissociation rates and diffusion coefficients, may be helpful. However, this approach may require specialized techniques to measure those parameters within the pericentriolar space. Alternatively, addition of other concepts may significantly improve these models. Introduction of the emerging concept of liquid–liquid phase separation may be one option (Woodruff et al., 2017). Furthermore, future experimental data may improve our mathematical models.

Our models principally assumed 12 slots in the CEP152 scaffold for Plk4. However, this cannot explain the nine-fold symmetric patterns of Plk4 that were observed following treatment with centrinone (Figure S6A). It is important to consider that treatment with centrinone may also affect cytosolic Plk4. Although centrinone is a powerful tool for the investigation of Plk4 functioning, alternative approaches may be necessary to precisely investigate the molecular dynamics in the nanoscopic space around centrioles. Another possible point to consider is that the CEP152 scaffold may be more flexible. Despite the apparent periodicity, the spatial patterns of CEP152 are somewhat less clear (Figure 3) than the sharp, discrete patterns of Plk4 (Figure 2). This could mean that the CEP152 scaffolds are more flexible and dynamic than we have supposed, and that the 12-fold rotational symmetry we have observed is merely their average form. Although our lateral inhibition model can simulate other numbers of slots in the scaffold (Figure S3), further studies are warranted to gain insight into the architecture of the CEP152 scaffold. Considering that the performance of STED microscopy may not be sufficient to reliably resolve such periodicities at scales less than 100 nm in our experimental conditions, future studies using electron microscopy or perhaps expansion microscopy (Chang et al., 2017; Chen et al., 2015) may provide more information regarding the scaffold for Plk4.

In conclusion, we propose here the first experimentally verifiable theory to explain Plk4’s formation of biased spatial patterns around centrioles, based on its intrinsic properties. In addition, we examined the involvement of this pattern formation in ensuring the exclusive provision of single sites for centriole duplication. This model sheds light on numerous experimental observations. However, further research based on both experimental data and further improvement of the present models may assist in reaching definitive conclusions.

## ACKNOWLEDGEMENTS

We gratefully acknowledge T. Fujiwara and iCeMS, Kyoto University, for technical support in STED microscopy; A. Kimura and D. Miyashiro for advice on mathematical modeling; M. Ohta, K. Watanabe, S. Yoshiba, and Y. Tsuchiya for technical support in the generation of the Plk4– mClover cell line and STED imaging of Plk4; and the members of the Kitagawa laboratory for technical support, discussion, and critical review of the manuscript. This work was supported by a Grant-in-Aid for Young Scientists (A) and Scientific Research (C) from the Ministry of Education, Science, Sports and Culture of Japan, by the Uehara Memorial Foundation, and by the Japan Prize Foundation.

## AUTHOR CONTRIBUTIONS

D.K. conceived the study. D.T. and D.K. designed the study and experiments. D.T. performed all the experiments and simulations. D.T. analyzed the data. D.T. and D.K. interpreted the data. D.T. and D.K. wrote the manuscript. S.Y. participated in the incubation of the core concept of the theory and reviewed the manuscript.

**Figure S1.**
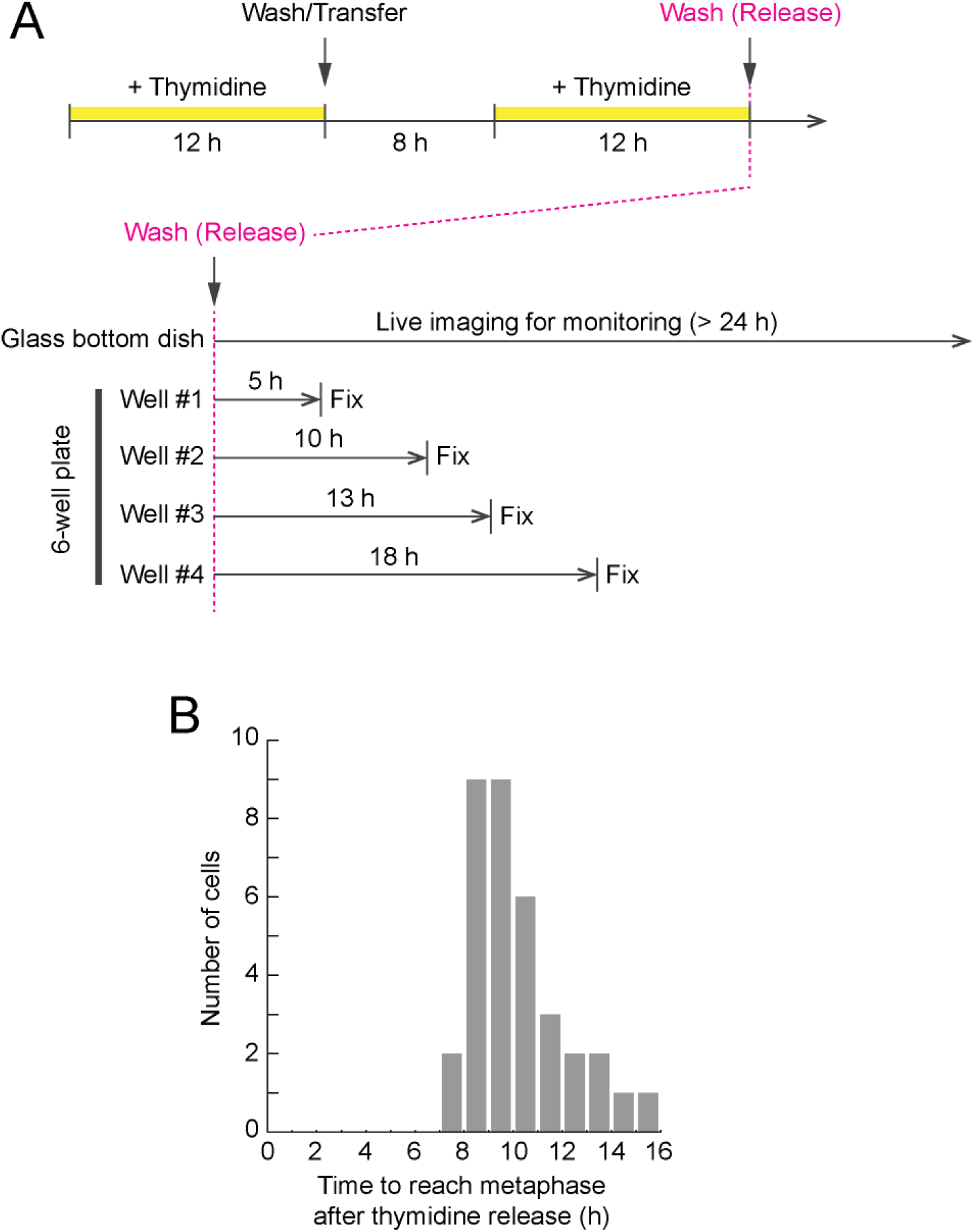
Time course of cell synchronization, for tracing changes in the spatial patterns of centriolar Plk4. (Related to Figure 1) (A) Time course of sample preparation after the synchronization of cells. Cells were arrested twice with thymidine. Following the second thymidine release, cells were fixed for immunofluorescence at the indicated time points, while the cell cycle was monitored via live-cell imaging. (B) The time taken by the monitored live cells to reach metaphase after thymidine release.

**Figure S2.**
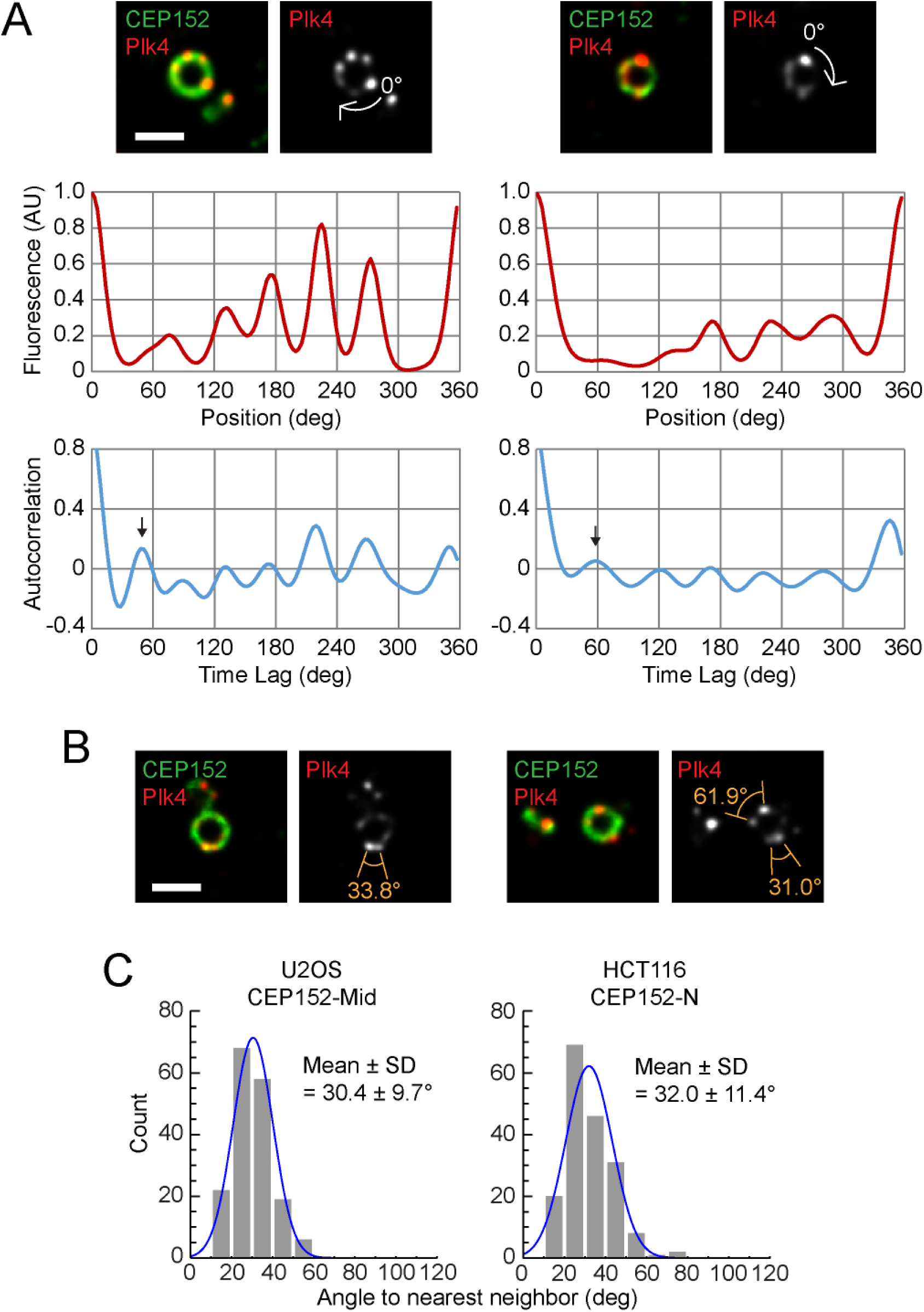
Periodicity of spatial patterns of centriolar Plk4 and CEP152. (Related to Figures 2 and 3) (A) Examples of autocorrelation analysis. Representative STED images (top), oval profiles (mid), and autocorrelation functions (bottom) are shown. The arrows in the autocorrelation graphs indicate the primary peaks representing their periods. Scale bar, 0.5 μm. (B) Representative STED images of Plk4 with foci at approximately 30° intervals. Scale bar, 0.5 μm. (C) Extended analyses of the periodicity of CEP152. Results of measurements similar to those presented in Figure 3 but using a different antibody (left) or another cell line (right) are shown. n = 19 and 20 cells, respectively.

**Figure S3.**
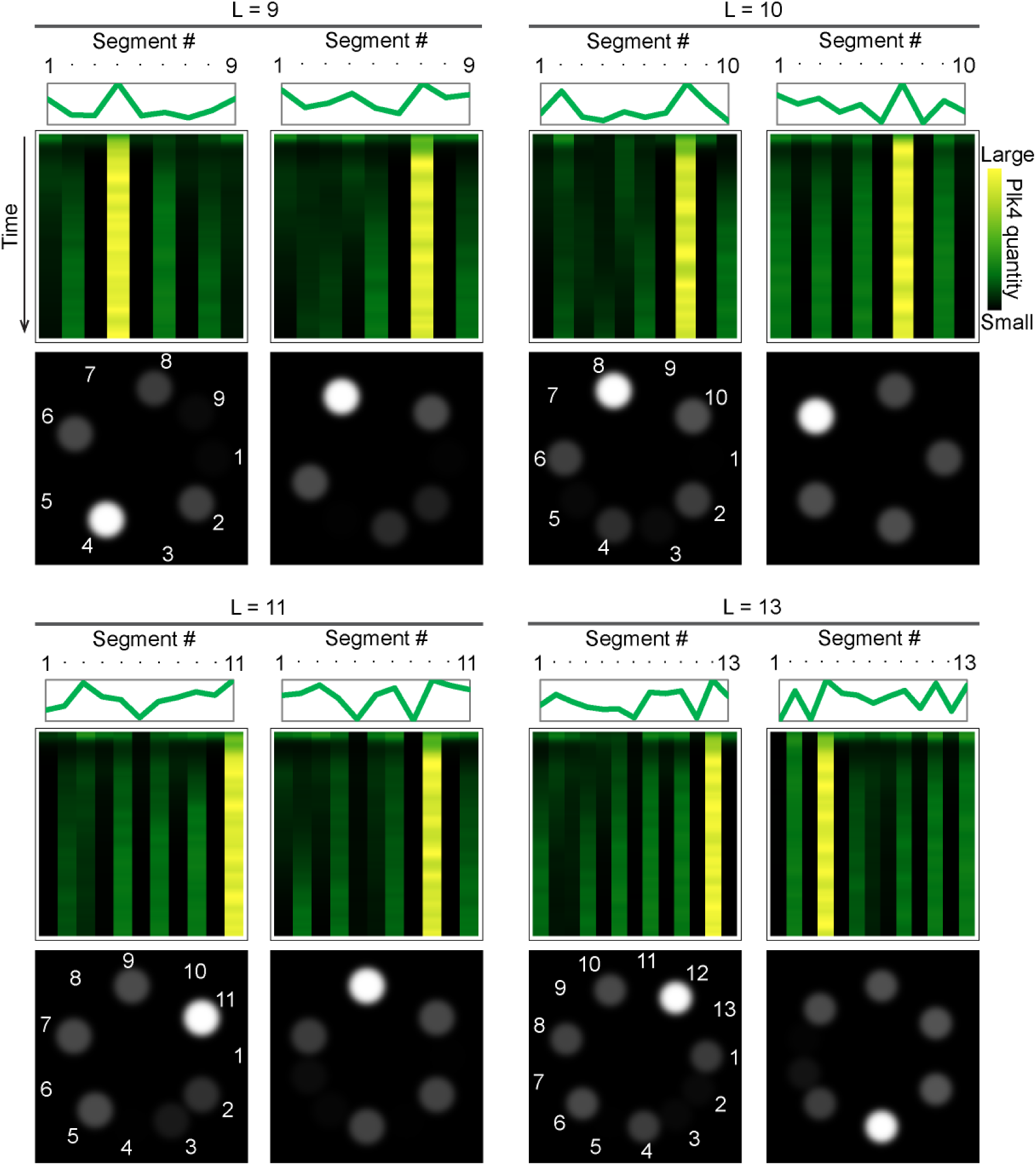
Simulations with variable scaffold numbers. (Related to Figure 5) Test of the dependency of the lateral inhibition model on the number of Plk4 slots. The number of slots (L) was set to 9, 10, 11, or 13 (in addition to 12, as shown in Figure 5). All simulations exhibited similar biased discrete patterns. Two examples are shown for each condition.

**Figure S4.**
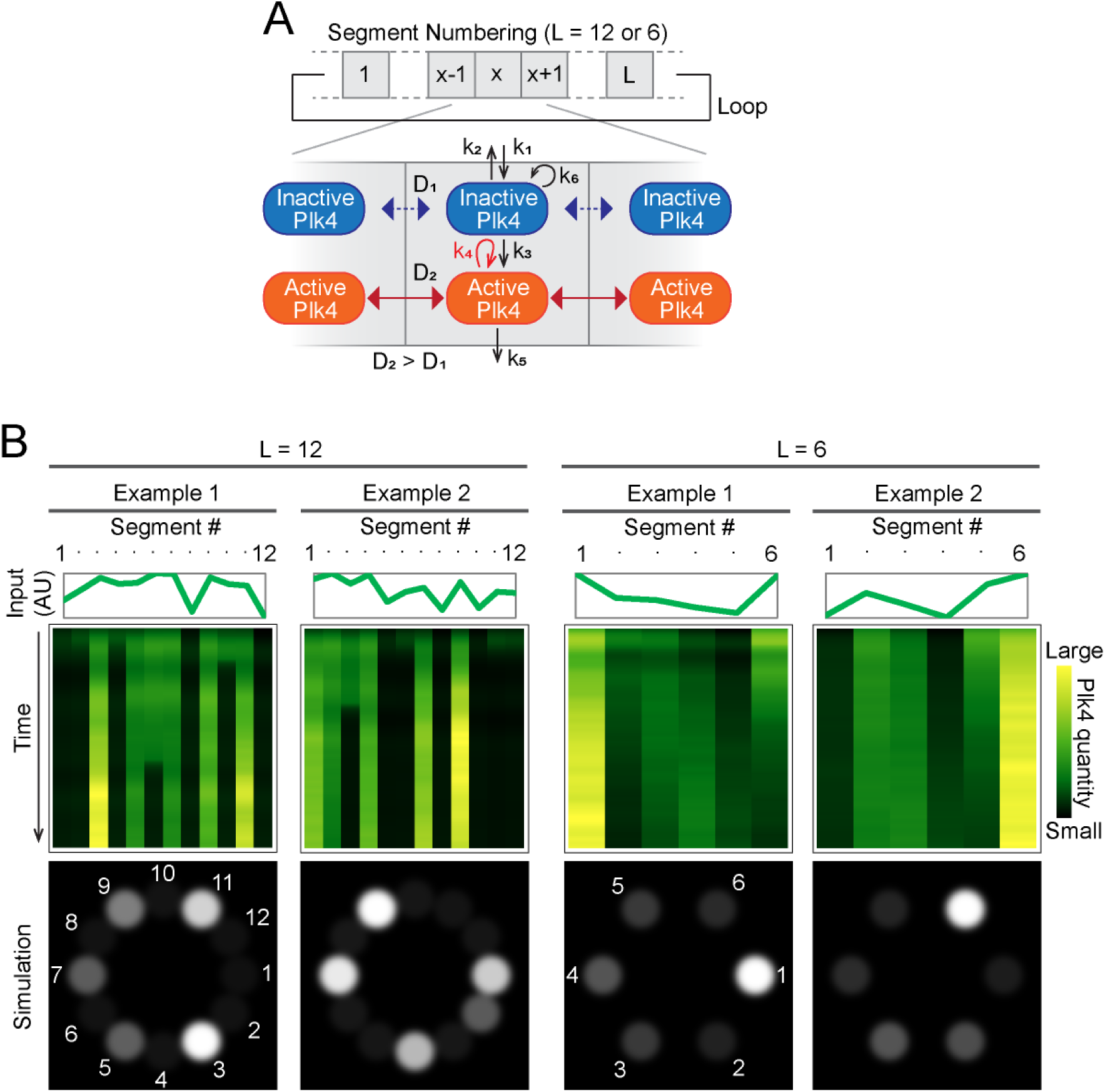
An alternative mathematical model for the pattern formation of Plk4. (Related to Figure 5) (A) Schematic drawing of the reaction–diffusion model. (B) Example results of the simulations.

**Figure S5.**
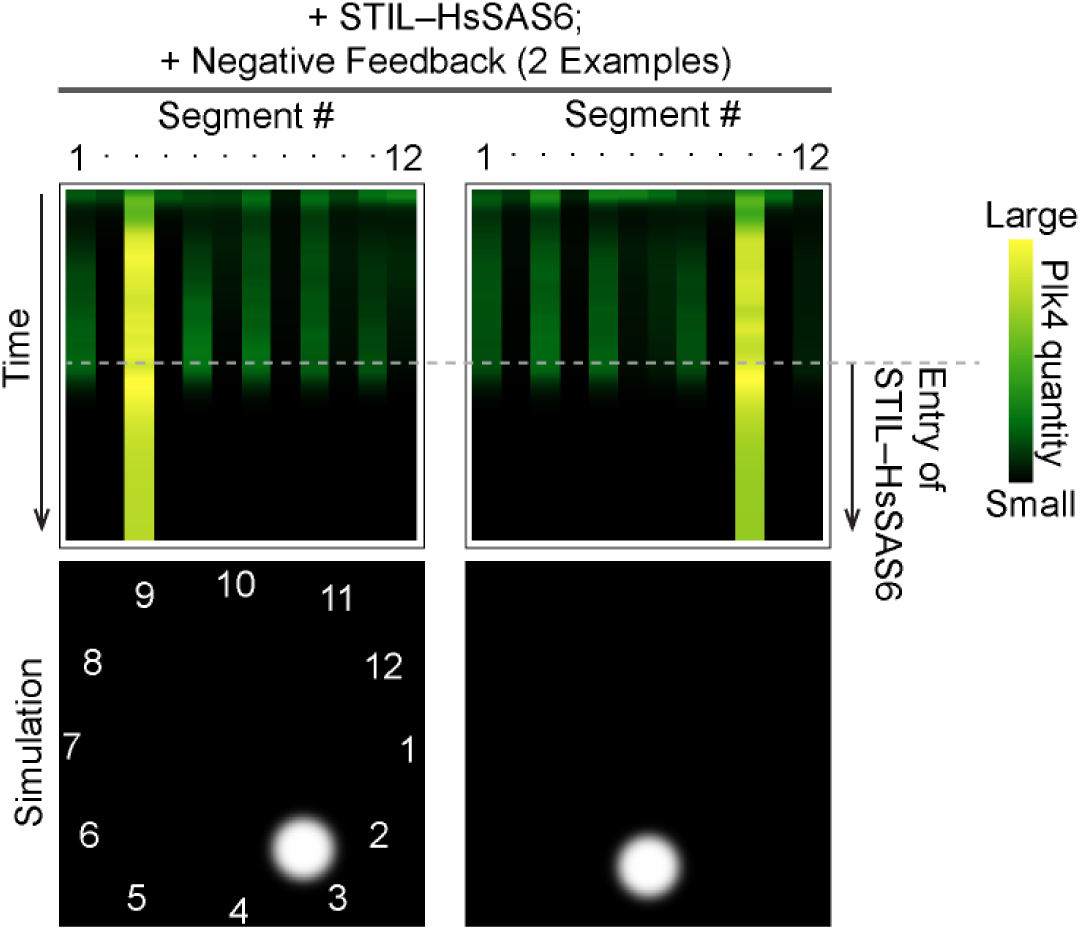
Simulations using the assumption of negative-feedback regulation. (Related to Figure 6) Representative simulation results using the assumption of negative-feedback regulation. At the mid-point of the time course, the STIL–HsSAS6 complex began to accumulate. It stabilized the dominant focus of Plk4, as in Figure 6, and affected the other segments negatively by promoting dissociation/degradation.

**Figure S6.**
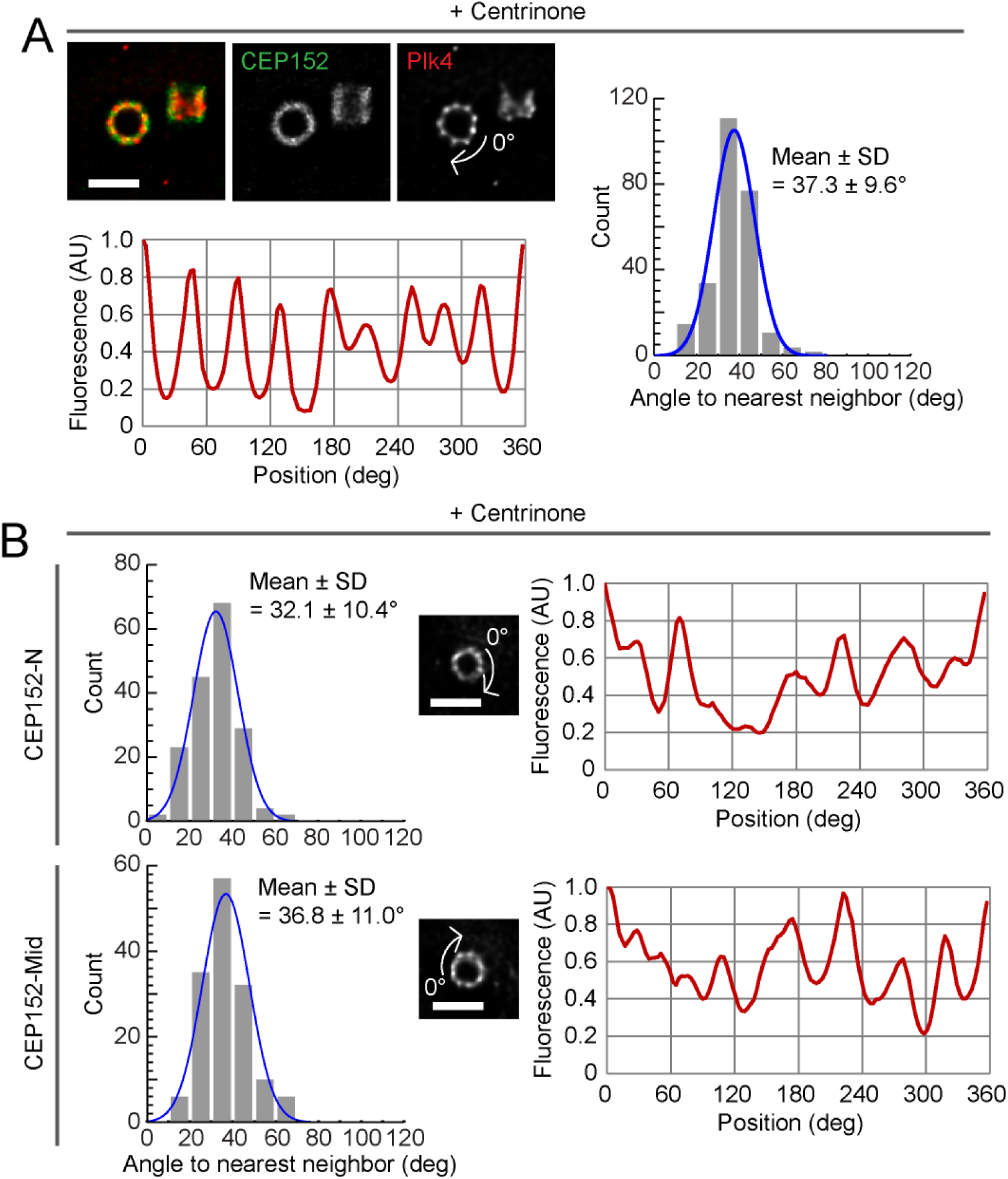
The effect of the inhibition of kinase activity on the pattern formation of Plk4. (Related to Figures 2 and 3) (A) STED pattern analysis of Plk4 in centrinone-treated cells. Data are displayed in a similar manner to those in Figures 2 and 3. Scale bar, 0.5 μm. n = 30 cells. (B) STED pattern analysis of CEP152 in centrinone-treated cells. Results using two different antibodies (CEP152-N/Mid) are shown. Scale bar, 0.5 μm. n = 29 and 18 cells, respectively.

## MATERIALS AND METHODS

### KEY RESOURCES

#### Antibodies

Plk4 (1:250; Merck, MABC544)

CEP152-N (1:1,000; Bethyl, A302-479A)

CEP152-Mid (1:1,000; Bethyl, A302-480A)

GFP (1:500; Invitrogen, A11120)

HA (1:500; Abcam, ab9110)

CP110 (1:1,000; Proteintech, 12780-1-AP)

Centrin (1:1,000; Merck, 04-1624)

#### Chemicals

Thymidine (Sigma, T1895)

Centrinone (MedChem Express, HY-18682).

## EXPERIMENTAL MODEL AND SUBJECT DETAILS

### Cell lines

HCT116 cells were cultured in McCoy’s 5A medium (GE Healthcare) supplied with 10% FBS, 1% glutamine, and 1% penicillin/streptomycin. U2OS cells were cultured in DMEM medium supplied with 10% FBS and 1% penicillin/streptomycin. For synchronization of the cell cycle, 2 mM thymidine were added to the medium, as indicated in Figure S1. For Plk4 inhibition, cells were incubated for 4 h in the presence of 200 nM centrinone prior to fixation.

Cells were transfected with DNA or siRNA using Lipofectamine 2000 or Lipofectamine RNAiMAX, according to the manufacturer’s instructions.

The HCT116 Plk4–mClover cell line was produced via CRISPR-Cas9 genome editing. The mClover sequence was inserted into the 3′ region of the Plk4 gene, and cell clones were selected using hygromycin. Cloned cells were genotyped using PCR, and the proper localization of expressed Plk4–mClover was verified via immunofluorescence. We were only able to obtain a monoallelic cell line, which we therefore used in this study.

### METHOD DETAILS

#### Immunofluorescence

Cells cultured on coverslips were fixed using cold methanol at −20 °C for 5 min. The fixed cells were washed three times with PBS and incubated in blocking buffer (1% BSA and 0.05% Triton X-100 in PBS) for 20 min at room temperature (RT). The cells were then incubated with primary antibodies in blocking buffer at 4 °C overnight, washed three times with PBS, and incubated with secondary antibodies in blocking buffer for 1 h at RT. The cells were stained with Hoechst 33258 (DOJINDO) in PBS for 5 min at RT, washed three times with PBS, and subsequently mounted with ProLong Gold (Thermo Fisher Scientific, #P36930).

#### Microscopy

For general observations, an upright epifluorescence microscope (Zeiss Axio Imager 2) with a 100× oil-immersion objective (N.A. 1.4) and an AxioCam HRm camera, or an inverted confocal microscope (Leica TCS SP8) equipped with a 63× oil-immersion objective (N.A. 1.4), was used. Z-stacked confocal images were obtained at 0.13 μm intervals.

For live-cell imaging, a spinning disc-based confocal microscope (Yokogawa, CV1000) equipped with a 60× oil-immersion objective (N.A. 1.35), a back-illuminated EMCCD camera, and a stage incubator supplied with 5% CO_2_ was used. Typically, 10–30 fields of view were recorded every 10 min for up to 30 h in a single experiment, and each field contained 25 z-slices with 1.3 μm intervals, subsequently max-projected using ImageJ software.

For STED observations, a STED system based on an inverted confocal microscope (Leica TCS SP8 STED) equipped with 592/660 STED laser lines and a 100× oil-immersion objective (N.A. 1.4) was used. The interval of optical sectioning was 0.18 μm.

The Huygens Essential or Professional image processing software was used for image deconvolution, and ImageJ software was used for image processing and analyses.

#### Mathematical modeling

To simulate centriole duplication, we constructed two mathematical models as follows. For convenience, here the models are termed the lateral inhibition (LI) and the reaction–diffusion (RD) models. Both models are based on the intrinsic properties of Plk4 that allow it to self-assemble and promote the dissociation/degradation of neighboring molecules following its activation. The units, including that used for time, are arbitrary (relative). As schematically shown in Figures 5A and S4A, there are only slight differences between the models. Specifically, the RD model more explicitly considers the lateral diffusion of Plk4 molecules within the periphery of the centriole. The periphery of the centriole (the scaffold of Plk4) is divided into segments (*x*), such that the last segment (*x* = L) is connected to the first (*x* = 1) to form a closed loop. L was typically assigned a value of 12. Plk4 can exist in the active (autophosphorylated) or the inactive form. In the LI model, the rate of change in the concentration of Plk4 at segment *x* and time *t* (I_*x*_(*t*) for the inactive form and A_*x*_(*t*) for the active form) is expressed using the following partial differential equations:

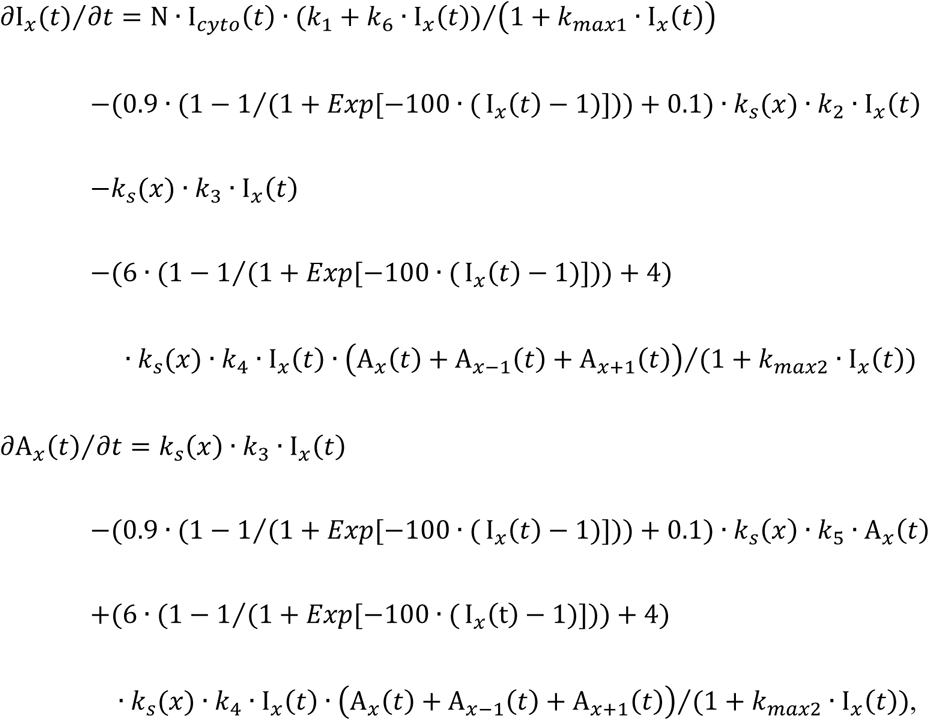

where N is a noise parameter, I_*cyto*_(*t*) is the cytosolic concentration of inactive Plk4, 1 + *k*_*max*1_ · I_*x*_(*t*) limits the centriolar influx of Plk4, and 0.9 · (1 − 1 (1 + *Exp*[−100 · (I_*x*_(*t*) −1)])) + 0.1 and 6 · (1 − 1 (1 + *Exp*[−100 · (I_*x*_(*t*) − 1)])) + 4 express the sigmoidal deceleration of Plk4 dissociation due to the decreased surface-to-volume ratio upon self-assembly. The parameter *k*_*s*_(*x*) decreases as STIL enters the segment and stabilizes Plk4. In the presence of cytosolic STIL, when the quantity of active Plk4 in segment *x*, A_*x*_(*t*), exceeds a given threshold (typically 0.5), STIL enters the segment. The initial values of the inactive form, I_*x*_(0), are provided as a list of random real numbers between 0 and 1, whereas those of the active form, A_*x*_(*t*), are set to 0. The basic parameter settings are listed in Table S1.

**Table S1.** Basic parameter settings for the LI model

| $k_1$ | $k_2$ | $k_3$ | $k_4$ | $k_5$ | $k_6$ | $k_{max1}$ | $k_{max2}$ | N | $I_{cyto}$ |
| --- | --- | --- | --- | --- | --- | --- | --- | --- | --- |
| 0.0001 | 0.01 | 0.008 | 0.01 | 0.1 | 0.04 | 0.02 | 0.06 | {0.9,1.1} | 0.8 |

The RD model explicitly includes diffusion formulae, and the concentration changes are similarly expressed as:

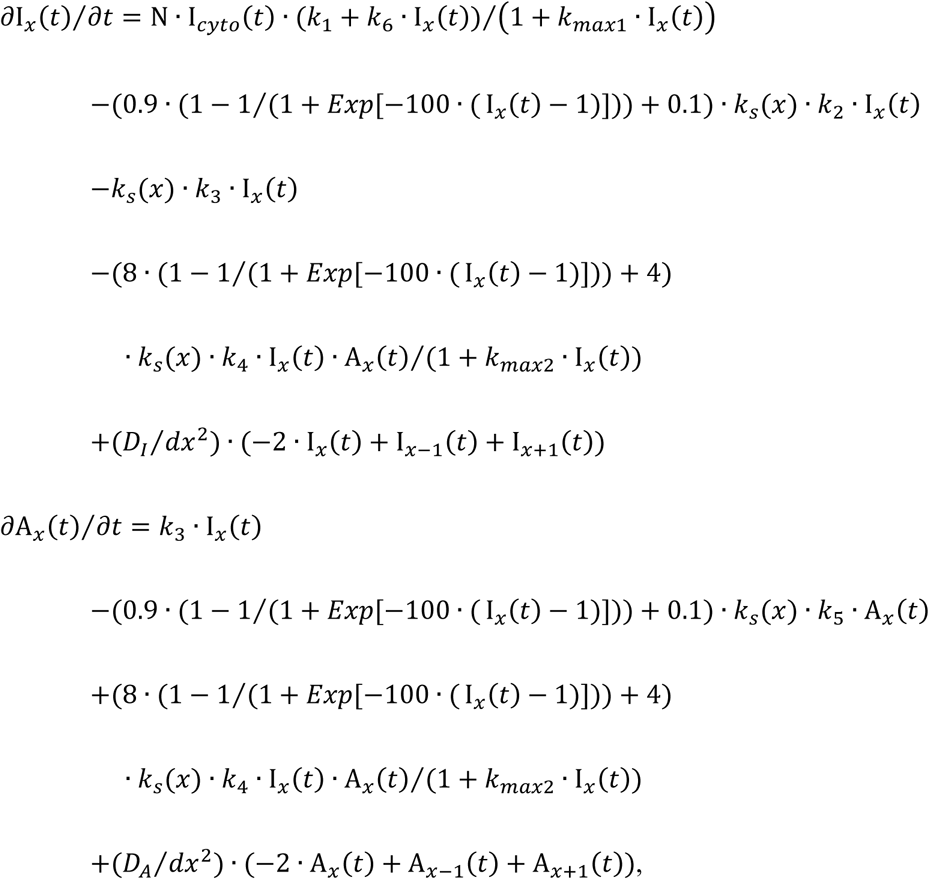

where *D_I_* and *D_A_* are the diffusion coefficients for the inactive and active forms of Plk4, respectively, and *dx* is the distance between each segment. The basic parameter settings are listed in Table S2.

**Table S2.** Basic parameter settings for the RD model

|  |  |  |  |  |  |  |  |  |  |
| --- | --- | --- | --- | --- | --- | --- | --- | --- | --- |
| $k_1$ | $k_2$ | $k_3$ | $k_4$ | $k_5$ | $k_6$ | $k_{\max 1}$ | $k_{\max 2}$ | N | $I_{\text{cyto}}$ |
| 0.0002 | 0.01 | 0.004 | 0.01 | 0.1 | 0.02 | 0.02 | 0.05 | {0.9,1.1} | 1 |
| $D_I$ | | $D_A$ | | $dx$ | | | | | |
| $1*10^{-6}$ | | $1*10^{-2}$ | | 0.01 | | | | | |

The simultaneous differential equations were numerically solved using our original Mathematica program. The program also generated the time-evolved simulation graphs and the simulated spatial distributions shown in Figures 5, 6, S3, and S4.

### QUANTIFICATION AND STATISTICAL ANALYSIS

All quantification and statistical analyses were performed using ImageJ, Mathematica, R, and Excel software. For live imaging, the fluorescence intensity of regions of interest of the same size was measured using ImageJ on max projection images and the fluorescence intensity of a no-cell region was used for background subtraction (Figure 1A). The oval profiles of Plk4 and CEP152 were measured using the Oval Profile Plot plugin with the “Along Oval” option, as shown in Figures 2B and 3. The number of sampling points was set to 64 for conventional confocal images or 128 for STED images, depending on image resolution. For the calculation of the ring-filling indices (Figure 1C), the profiles were exported to and processed in Excel. The box-and-whisker plots in Figure 1D were generated using R. The profiles were exported to Mathematica to generate the 2D array plot shown in Figure 2C. Using our Mathematica programs, the peaks in the profiles were detected by calculating the local maxima, and the angles (distances) between nearest-neighbor pairs of foci were obtained, as shown in Figures 2D and 3. In addition, we developed a Mathematica program to obtain autocorrelation functions (Figure S2A). The original data were zero-padded for the calculations.

## REFERENCES

Arquint, C., and Nigg, E.A. (2014). STIL microcephaly mutations interfere with APC/C-mediated degradation and cause centriole amplification. Curr. Biol. 24, 351–360.

Arquint, C., Sonnen, K.F., Stierhof, Y.-D., and Nigg, E.A. (2012). Cell-cycle-regulated expression of STIL controls centriole number in human cells. J. Cell Sci. 125, 1342–1352.

Arquint, C., Gabryjonczyk, A.M., Imseng, S., Böhm, R., Sauer, E., Hiller, S., Nigg, E.A., and Maier, T. (2015). STIL binding to Polo-box 3 of PLK4 regulates centriole duplication. Elife 4, 1–22.

Banterle, N., and Gönczy, P. (2017). Centriole Biogenesis: From Identifying the Characters to Understanding the Plot. Annu. Rev. Cell Dev. Biol. 33, 23–49.

Barad, O., Hornstein, E., and Barkai, N. (2011). Robust selection of sensory organ precursors by the Notch-Delta pathway. Curr. Opin. Cell Biol. 23, 663–667.

Bettencourt-Dias, M., Rodrigues-Martins, A., Carpenter, L., Riparbelli, M., Lehmann, L., Gatt, M.K., Carmo, N., Balloux, F., Callaini, G., and Glover, D.M. (2005). SAK/PLK4 is required for centriole duplication and flagella development. Curr. Biol. 15, 2199–2207.

Chang, J.-B., Chen, F., Yoon, Y.-G., Jung, E.E., Babcock, H., Kang, J.S., Asano, S., Suk, H.-J., Pak, N., Tillberg, P.W., et al. (2017). Iterative expansion microscopy. Nat. Methods 14, 593–599.

Chen, F., Tillberg, P.W., and Boyden, E.S. (2015). Optical imaging. Expansion microscopy. Science 347, 543–548.

Chen, Q., Shi, J., Tao, Y., and Zernicka-Goetz, M. (2018). Tracing the origin of heterogeneity and symmetry breaking in the early mammalian embryo. Nat. Commun. 9, 1819.

Cizmecioglu, O., Arnold, M., Bahtz, R., Settele, F., Ehret, L., Haselmann-Weiss, U., Antony, C., and Hoffmann, I. (2010). Cep152 acts as a scaffold for recruitment of Plk4 and CPAP to the centrosome. J. Cell Biol. 191, 731–739.

Dzhindzhev, N.S., Yu, Q.D., Weiskopf, K., Tzolovsky, G., Cunha-Ferreira, I., Riparbelli, M., Rodrigues-Martins, A., Bettencourt-Dias, M., Callaini, G., and Glover, D.M. (2010). Asterless is a scaffold for the onset of centriole assembly. Nature 467, 714–718.

Dzhindzhev, N.S., Tzolovsky, G., Lipinszki, Z., Schneider, S., Lattao, R., Fu, J., Debski, J., Dadlez, M., and Glover, D.M. (2014). Plk4 phosphorylates Ana2 to trigger Sas6 recruitment and procentriole formation. Curr. Biol. 24, 2526–2532.

Dzhindzhev, N.S., Tzolovsky, G., Lipinszki, Z., Abdelaziz, M., Debski, J., Dadlez, M., and Glover, D.M. (2017). Two-step phosphorylation of Ana2 by Plk4 is required for the sequential loading of Ana2 and Sas6 to initiate procentriole formation. Open Biol. 7.

Goryachev, A.B., and Leda, M. (2017). Many roads to symmetry breaking: molecular mechanisms and theoretical models of yeast cell polarity. Mol. Biol. Cell 28, 370–380.

Habedanck, R., Stierhof, Y.-D., Wilkinson, C.J., and Nigg, E.A. (2005). The Polo kinase Plk4 functions in centriole duplication. Nat. Cell Biol. 7, 1140–1146.

Halatek, J., Brauns, F., and Frey, E. (2018). Self-organization principles of intracellular pattern formation. Philos. Trans. R. Soc. B Biol. Sci. 373, 20170107.

Hatch, E.M., Kulukian, A., Holland, A.J., Cleveland, D.W., and Stearns, T. (2010). Cep152 interacts with Plk4 and is required for centriole duplication. J. Cell Biol. 191, 721–729.

Izraeli, S., Colaizzo-Anas, T., Bertness, V.L., Mani, K., Aplan, P.D., and Kirsch, I.R. (1997). Expression of the SIL gene is correlated with growth induction and cellular proliferation. Cell Growth Differ. 8, 1171–1179.

Jana, S.C., Mendonça, S., Machado, P., Werner, S., Rocha, J., Pereira, A., Maiato, H., and Bettencourt-Dias, M. (2018). Differential regulation of transition zone and centriole proteins contributes to ciliary base diversity. Nat. Cell Biol. 20, 928–941.

Kim, E.J.Y., Korotkevich, E., and Hiiragi, T. (2018). Coordination of Cell Polarity, Mechanics and Fate in Tissue Self-organization. Trends Cell Biol. 28, 541–550.

Kleylein-Sohn, J., Westendorf, J., Le Clech, M., Habedanck, R., Stierhof, Y.-D., and Nigg, E.A. (2007). Plk4-induced centriole biogenesis in human cells. Dev. Cell 13, 190–202.

Kondo, S., and Miura, T. (2010). Reaction-Diffusion Model as a Framework for Understanding Biological Pattern Formation. Science. 329, 1616–1620.

Liao, B.-K., and Oates, A.C. (2017). Delta-Notch signalling in segmentation. Arthropod Struct. Dev. 46, 429–447.

McLamarrah, T.A., Buster, D.W., Galletta, B.J., Boese, C.J., Ryniawec, J.M., Hollingsworth, N.A., Byrnes, A.E., Brownlee, C.W., Slep, K.C., Rusan, N.M., et al. (2018). An ordered pattern of Ana2 phosphorylation by Plk4 is required for centriole assembly. J. Cell Biol. 217, 1217–1231.

Moyer, T.C., Clutario, K.M., Lambrus, B.G., Daggubati, V., and Holland, A.J. (2015). Binding of STIL to Plk4 activates kinase activity to promote centriole assembly. J. Cell Biol. 209, 863–878.

Nakamura, T., Mine, N., Nakaguchi, E., Mochizuki, A., Yamamoto, M., Yashiro, K., Meno, C., and Hamada, H. (2006). Generation of robust left-right asymmetry in the mouse embryo requires a self-enhancement and lateral-inhibition system. Dev. Cell 11, 495–504.

Nigg, E.A., and Holland, A.J. (2018). Once and only once: mechanisms of centriole duplication and their deregulation in disease. Nat. Rev. Mol. Cell Biol. 19, 297–312.

Ohta, M., Ashikawa, T., Nozaki, Y., Kozuka-Hata, H., Goto, H., Inagaki, M., Oyama, M., and Kitagawa, D. (2014). Direct interaction of Plk4 with STIL ensures formation of a single procentriole per parental centriole. Nat. Commun. 5, 5267.

Ohta, M., Watanabe, K., Ashikawa, T., Nozaki, Y., Yoshiba, S., Kimura, A., and Kitagawa, D. (2018). Bimodal Binding of STIL to Plk4 Controls Proper Centriole Copy Number. Cell Rep. 23, 3160–3169.e4.

Saha, S., Nagy, T.L., and Weiner, O.D. (2018). Joining forces: crosstalk between biochemical signalling and physical forces orchestrates cellular polarity and dynamics. Philos. Trans. R. Soc. B Biol. Sci. 373, 20170145.

Shi, X., Garcia, G., Van De Weghe, J.C., McGorty, R., Pazour, G.J., Doherty, D., Huang, B., and Reiter, J.F. (2017). Super-resolution microscopy reveals that disruption of ciliary transition-zone architecture causes Joubert syndrome. Nat. Cell Biol. 19, 1178–1188.

Strnad, P., Leidel, S., Vinogradova, T., Euteneuer, U., Khodjakov, A., and Gönczy, P. (2007). Regulated HsSAS-6 Levels Ensure Formation of a Single Procentriole per Centriole during the Centrosome Duplication Cycle. Dev. Cell 13, 203–213.

Sych, T., Mély, Y., and Römer, W. (2018). Lipid self-assembly and lectin-induced reorganization of the plasma membrane. Philos. Trans. R. Soc. B Biol. Sci. 373, 20170117.

Tang, C.-J.C., Lin, S.-Y., Hsu, W.-B., Lin, Y.-N., Wu, C.-T., Lin, Y.-C., Chang, C.-W., Wu, K.-S., and Tang, T.K. (2011). The human microcephaly protein STIL interacts with CPAP and is required for procentriole formation. EMBO J. 30, 4790–4804.

Vulprecht, J., David, A., Tibelius, A., Castiel, A., Konotop, G., Liu, F., Bestvater, F., Raab, M.S., Zentgraf, H., Izraeli, S., et al. (2012). STIL is required for centriole duplication in human cells. J. Cell Sci. 125, 1353–1362.

Wheeler, R.J., and Hyman, A.A. (2018). Controlling compartmentalization by non-membrane-bound organelles. Philos. Trans. R. Soc. B Biol. Sci. 373, 20170193.

Wong, Y.L., Anzola, J.V, Davis, R.L., Yoon, M., Motamedi, A., Kroll, A., Seo, C.P., Hsia, J.E., Kim, S.K., Mitchell, J.W., et al. (2015). Cell biology. Reversible centriole depletion with an inhibitor of Polo-like kinase 4. Science 348, 1155–1160.

Woodruff, J.B., Ferreira Gomes, B., Widlund, P.O., Mahamid, J., Honigmann, A., and Hyman, A.A. (2017). The Centrosome Is a Selective Condensate that Nucleates Microtubules by Concentrating Tubulin. Cell 169, 1066–1077.e10.

Yamamoto, S., and Kitagawa, D. (2018). Self-organization of Plk4 regulates symmetry breaking in centriole duplication. BioRxiv. doi: http://dx.doi.org/10.1101/313635.

Yang, T.T., Chong, W.M., Wang, W.-J., Mazo, G., Tanos, B., Chen, Z., Tran, T.M.N., Chen, Y.-D., Weng, R.R., Huang, C.-E., et al. (2018). Super-resolution architecture of mammalian centriole distal appendages reveals distinct blade and matrix functional components. Nat. Commun. 9, 2023.

